# α-Synuclein Activates the PI3K/AKT Pathway to Drive Lipid Droplets Accumulation: Implications for Parkinson’s Disease

**DOI:** 10.1101/2025.03.27.645689

**Authors:** Suaad Abd Elhadi, Jenna Congdon-Loeffler, Eva Harrosch, Dima Khatib, Lama Naamneh, Meir Schechter, Alexandra Eliassaf, Ori Shalev, Ronit Sharon

## Abstract

Growing evidence supports a metabolic component in Parkinson’s disease (PD). α- Synuclein (α-Syn), a protein central to the onset and progression of PD, facilitates the accumulation of neuronal lipid droplets, which are implicated in disease pathology. We report that AKT is hyper-phosphorylated in PD brains and show that α-Syn enhances p110α activity by facilitating palmitoylated Ras localization to the plasma membrane, driving lipid droplet accumulation through PI3K/AKT/mTOR and PPARγ activation.

Phosphorylation of α-Syn at Ser129 correlates positively with the localization of Ras to the membrane fraction and with accumulation of lipid droplets. In-vivo treatment of young, asymptomatic α-Syn^A53T^ transgenic mice with GDC-0084 (paxalisib), a blood-brain barrier-permeable PI3K inhibitor, restored healthy AKT activity levels, reduced levels of PSer129 and α-Syn oligomers, decreased neuronal lipid droplet accumulation, and promoted lysosomal clustering. These findings establish a role for α-Syn in p110α activation during early, asymptomatic stages of the disease and highlight the therapeutic potential of PI3K inhibition as a disease-modifying strategy.

**Highlight:** α-Syn activates PI3K/AKT by promoting the localization of palmitoylated Ras to membranes, linking metabolic dysfunction to PD.

## Introduction

Parkinson’s disease (PD) is a heterogeneous disorder with an unclear etiology(*1*). α- Synuclein (α-Syn) is central to PD genetics, pathology, and clinical progression(*2*). Growing evidence implicates metabolic dysfunction in PD, with genetic studies highlighting the role of energy and lipid metabolism(*3*). Additionally, metabolic disorders correlate with PD onset and severity, and metabolic drugs show promise in recent clinical trials(*4*).

Lipid droplets (LDs) are dynamic lipid storage organelles linked to cell energy metabolism. They play protective roles in stress conditions (*5*) and are implicated in the pathology of PD(*6*). Increases in LDs has been observed in models overexpressing α-Syn including, yeast cells(*7*), non-neuronal cell lines(*8*) and neurons(*9, 10*). LD accumulation has been associated with α-Syn activity(*10, 11*) and its toxicity(*9, 10, 12*).

Phosphoinositide 3-kinases (PI3Ks) are a family of lipid-kinases that can be classified into three classes based on their structural organization and functional characteristics(*13*).

Class I PI3Ks, in particular, are receptor-activated enzymes that play a crucial role in cell signaling by synthesizing phosphatidylinositol (PI) 3,4,5-trisphosphate (PI3,4,5P_3_) at the plasma membrane(*14*). These enzymes are heterodimeric, composed of a catalytic subunit (p110α, p110β, p110γ or p110δ) and a regulatory subunit(*15*). Among the four catalytic isoforms, p110α, encoded by PIK3CA, is frequently implicated in the development and progression of various human diseases, most notably in malignancies, due to its common mutations(*16*). The lipid kinase activity of p110α is regulated by the p85 regulatory subunit and by activated Ras proteins which bind directly to the N-terminal RAS-binding domain (RBD) of p110α (*17*). In contrast to PIK3CA, transforming mutations in the PIK3CB gene encoding p110β are less common, however, the gene is ubiquitously expressed, and was shown to be modulated in response to PIK3CA inhibition(*18*).

The PI3K/AKT/mTOR pathway is a central intracellular cascade that regulates cell survival, growth and metabolism. This pathway is activated in response to external stimuli, including growth factors and hormones that bind to specific membrane receptors (*19*). Dysregulation of the PI3K/AKT/mTOR pathway have been associated with the pathogenesis of PD(*20, 21*), primarily due to its key roles in autophagy and mitophagy(*22*).

AKT is activated by PDK1 and mTORC2, and subsequently activates mTORC1(*23*). mTOR complexes play distinct roles in LDs accumulation. While mTORC1 promotes de-novo lipid synthesis via sterol regulatory element-binding proteins (SREBPs) (*23, 24*), mTORC2 facilitates hepatic LD accumulation via activation of ATP citrate lyase (ACLY)(*25, 26*). In endothelial cells, both mTORC1 and mTORC2 were shown to regulate ACLY activation(*27*). In addition, both complexes are implicated in the activation of PPARγ, which is central to the regulation of lipogenesis (*28, 29*).

We report that α-Syn facilitates the activation of the p110α subunit of PI3K by promoting the plasma membrane localization of palmitoylated Ras proteins. This activation triggers downstream mTOR/PPARγ signaling, driving α-Syn-mediated neuronal lipid droplet accumulation. Furthermore, dynamic changes in PSer129 α-Syn levels, which strongly correlate with PI3K/AKT activation, highlight a pathogenic feedback between pathway hyperactivation and α-Syn activity/toxicity. Notably, the observed hyper-phosphorylation of AKT in PD brains supports the rationale for treating young and apparently healthy α-

Syn^A53T^ transgenic mice with GDC-0084 (paxalisib), a blood-brain barrier-permeable PI3K inhibitor. The in-vivo results highlight the therapeutic potential of targeting the PI3K/AKT pathway early in PD.

## Results

### The PI3K/AKT pathway is activated in PD brains and models consisting of α-Syn expression

To find out whether the canonical PI3K/AKT pathway contributes to PD pathogenesis, we assessed AKT phosphorylation in human PD and control brains. Frozen caudate (striatum) tissues were analyzed, with phosphorylated AKT levels normalized to total AKT levels within the same samples. The results revealed hyperactivation of the pathway, with elevated levels of phosphorylated AKT at Thr 308 and a significantly higher levels of phosphorylated AKT at Ser 473 in PD brains compared to age- and gender-matched, non-demented, control brains (n=12 per group; Fig. 1A,B).

**Fig 1.**
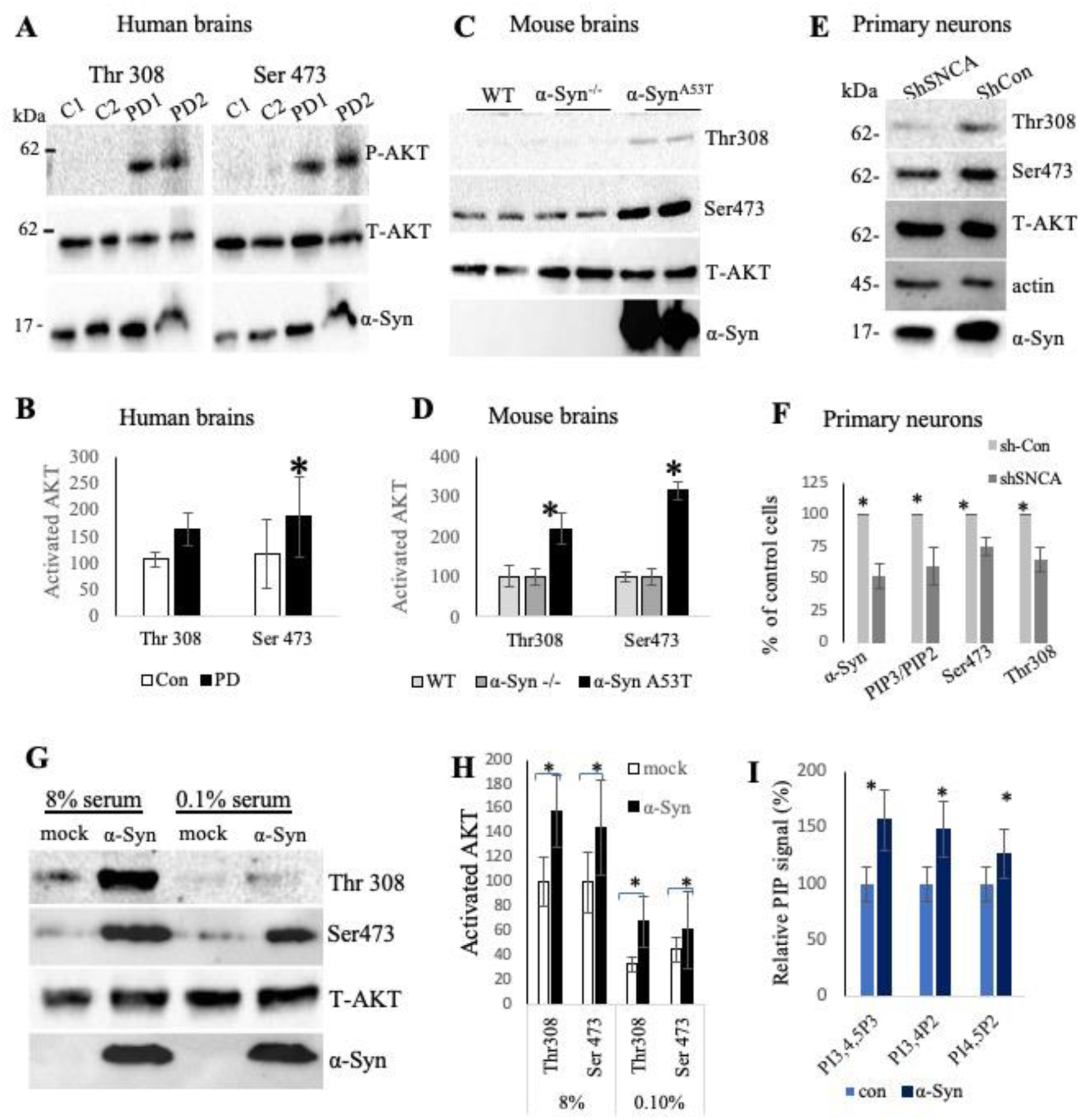
The PI3K/AKT pathway is activated in human PD brains and disease models involving α-Syn. **(A)** Western blotting of tissue homogenates from human brains containing the caudate. Immunoreacted with antibodies against phospho- AKT at Thr308 or Ser473, total AKT (T-AKT) and α-Syn. A representative blot of n=10. **(B)** Graph showing phospho AKT out of total AKT, presented as percent of control brains, set at 100%. Mean ± SE of n=12 brains in each group, analyzed in three different dates, *,p < 0.05, ttest. **(C)** Western blotting of whole mouse brain homogenates from α-Syn^A53T^, α-Syn^−/−^ (C57BL/6JOlaHsd) or WT (C57BL/6JRccHsd). Immunoreacted against Thr308 and Ser473, and total AKT. A representative image out of n=4. **(D)** Graph showing Mean ± SD of 4-6 mice in each genotype, including males and females. * P< 0.05, ANOVA. **(E)** Primary cortical neurons dissociated from α-Syn^A53T^ mouse brain, infected with shSNCA (Mission, Sigma) or a scrambled shRNA (shCon) and analyzed by Western blotting at 10 DIV. Immunoblot reacted with antibodies for α-Syn, Thr308, Ser473 and total-AKT levels. Actin levels for loading control. **(F)** Mean± SD of n=2 independent experiments as in (E). shCon is set at 100% for each variable. PI3,4,5P_3_ (PIP_3_) and PI4,5P_2_ (PIP_2_) levels determined by FACS following incubation of cells with specific antibodies (Echelon Biosciences).*, p<0.01, ttest. (**G)** HepG2 cells infected to express α-Syn or mock vector for 48 hours and conditioned either in serum supplemented EMEM (8%) or serum-deprived EMEM (0.1% serum) before cell lysis and analysis by Western blotting. **(H)** Mean± SD of n=3 experiments as in (G). *, p< 0.05, ttest. **(I)** HepG2 cells infected to express either a mRuby-P2A-α-Syn plasmid (α-Syn) or a mock-P2AmRuby plasmid (con) for 72 hours. PIP levels determined by FACS, after overnight starvation (0.1% FBS in DMEM) followed by serum replenishment for one hour, using anti-PI(4,5)P_2_, anti-PI(3,4,5)P_3_, and anti-PI(3,4)P_2_ abs. Mean ± SD of > 25000 gated cells, n=4 independent experiments in HepG2 cells, with the control (con) set at 100% for each variable. *, p<0.1, ttest.

PD is a multifactorial syndrome(*1*) and α-Syn is critically implicated in its pathogenesis(*2*). To investigate whether α-Syn expression is associated with PI3K/AKT activation, we analysed whole brains of α-Syn^A53T^ tg mice and two control mouse genotypes, WT (C57BL*/*6J*-*RccHsd) or α-Syn^-/-^ (C57BL/6JolaHsd). In young (3–4 months old), non-symptomatic α-Syn^A53T^ mice, we observed significantly elevated levels of phosphorylated AKT at both Thr308 and Ser473 compared to the age-matched control mice (Fig. 1C,D). Notably, no differences were detected between the two control mouse lines. In primary cortical neurons dissociated from WT C57BL/6 mouse brains, silencing endogenous α-Syn expression with shSNCA resulted in lower AKT phosphorylation levels compared to neurons expressing a scrambled shRNA (Fig. 1E,F). Consistent with the reduced AKT activation with α-Syn silencing, we also observed a lower PIP_3_ to PI4,5P_2_ ratio, indicating PI3K inhibition (Fig. 1F).

Furthermore, expressing α-Syn for 48 hours in HepG2 cells, which are known for their energy and lipid metabolic gene expression, resulted in a significant increase in AKT phosphorylation at Thr308 and Ser473. Serum starvation (0.1% FBS-supplemented EMEM) for 16 hours lowered overall AKT phosphorylation; however, α-Syn-expressing cells still maintained higher AKT phosphorylation levels over the control cells(Fig. 1G,H). Consistent with PI3K/AKT activation in the α-Syn-expressing HepG2 cells, we observed increased levels of PIPs, including PI3,4,5P_3_, PI4,5P_2_, and PI3,4P_2_, as measured by FACS (Fig. 1I).

### α-Syn activates p110α (PIK3CA) and P110β (PIK3CB)

To examine the specific role of p110α and/or p110β in α-Syn-induced AKT phosphorylation, we tested the effects of their respective inhibitors, PIK75 and TGX-221. HepG2 cells expressing either α-Syn or a mock vector were treated with the specified inhibitors for three hours before collection and analysis by Western blotting (Fig. 2A,B). The results showed a decrease in AKT phosphorylation at Thr308 and Ser473 with either inhibitor alone, and a greater reduction when both inhibitors were used in combination (Fig. 2A,B). Importantly, the differences in AKT phosphorylation between α-Syn-expressing and control cells were eliminated with the inhibitors, suggesting that α-Syn-induced hyperphosphorylation of AKT is mediated through PI3K activation.

**Fig. 2.**
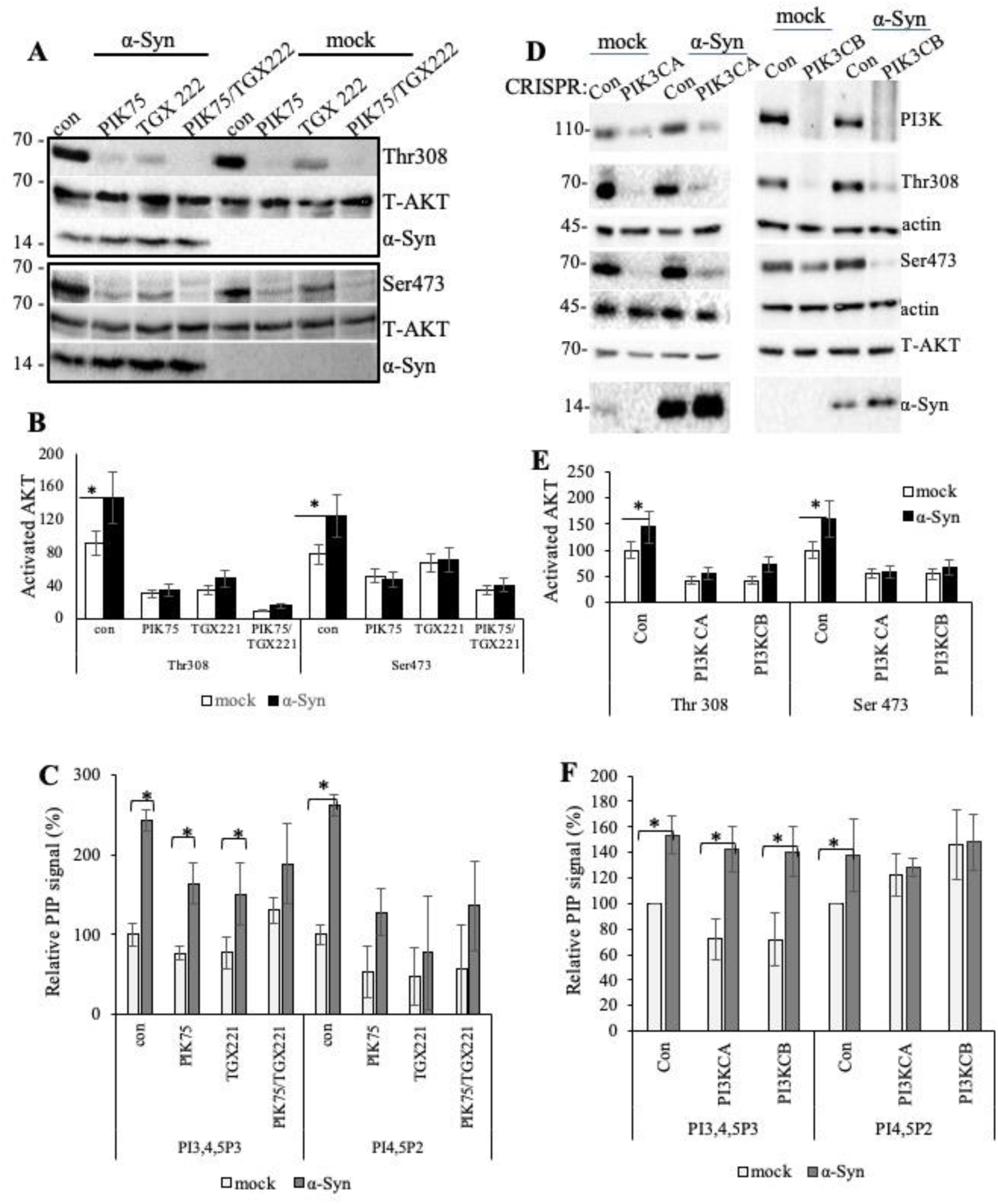
α-Syn activates PI3K p110α and p110β. **(A)** HepG2 cells were infected to express α-Syn or a mock vector. Cells were serum-starved overnight in EMEM containing 0.1% serum. Subsequently, cells were conditioned for additional 3 hours in media supplemented with 8% serum, together with the indicated inhibitors [PIK75 (100 nM), TGX 221 (100 nM), or a combination of both]. Cells were harvested 72 hours post-infection and analyzed via Western blotting. A representative blot from n=3 independent experiments. **(B)** Bar graph showing mean±SD of n=4 trials, *, p< 0.01, ANOVA. **(C)** Detection of PI3,4,5P_3_ and PI4,5P_2_ in cells treated as in (A) via FACS and specific anti-PIP antibodies. Graph showing Mean± SD of n=5 independent experiments. Control cells set at 100%. *, p< 0.01, ANOVA. **(D)** HepG2 cells were co-infected with a mock vector or α-Syn together with a CRISPR/CAS vector to silence either PI3KCA (p110α) or PI3KCB (p110β), or a control gRNA (Con). 72 hours post infection, cells were processed for the detection of the indicated proteins. A representative Western blot out of n=3 experiments. **(E)** Graph showing levels of Thr308 and Ser 473 normalized to total AKT as in (D). Mean±SD of n=3 repeats. Control cells set at 100%. *, p< 0.05, ANOVA. **(F)** Detection of PI3,4,5P_3_ and PI4,5P_2_ following CRISPR/CAS silencing of PI3KCA or PI3KCB in cells, via FACS. Mean±SD of n=4 repeats. *, p< 0.05, ANOVA.

Silencing either PIK3CA or PIK3CB in HepG2 cells was achieved via CRISPR/Cas with target-specific gRNAs. Cells were co-transfected with either α-Syn or a mock vector alongside the CRISPR/Cas construct. Controls included a scrambled gRNA (Con). The results showed a general reduction in phospho-AKT levels in cells where PIK3CA or PIK3CB expression was silenced. Notably, α-Syn-induced increases in AKT phosphorylation were abolished following CRISPR/Cas-mediated silencing of either PIK3CA or PIK3CB (Fig. 2D,E).

Consistent with the decreased AKT phosphorylation, treatment with the specific inhibitors resulted in lower PIP_3_ levels. However, α-Syn expression still led to higher PIP_3_ levels compared to those observed in mock-expressing cells (Fig. 2C). Similarly, CRISPR/Cas-mediated downregulation of either PIK3CA or PIK3CB resulted in lower PIP_3_ levels in control cells (Fig. 2F). However, in α-Syn-expressing cells, PIP_3_ levels were not affected with CRISPR/Cas (Fig. 2F). This suggests that alternative mechanisms may be activated to maintain PIP_3_ levels in α-Syn-expressing cells upon downregulation of PIK3CA or PIK3CB expression. Such compensatory mechanisms appear more evident following 72- hour CRISPR/Cas intervention compared to the shorter three-hour pharmacological inhibition.

### α-Syn enhances the accumulation of LDs in neuronal cells in a mechanism dependent of PI3K/AKT activation

Since both the PI3K/AKT pathway and α-Syn are linked to LD accumulation (*7–11, 30–32*) we hypothesized that α-Syn promotes LDs via this pathway. To test this, α-Syn^-/-^ cortical neurons were infected with mock or α-Syn-expressing vectors. Neurons were treated with GDC-0084, a PI3K inhibitor or vehicle and stained for LDs using Bodipy 493/503, a dye that specifically labels neutral lipids within LDs. The results show accumulation of LDs in α-Syn-positive neurons but not in the mock-infected neurons (Fig. 3A,B). Importantly, a significantly lower Bodipy 493/503 signal is detected in the α-Syn-positive neurons treated with GDC-0084 (Fig. 3A,B). In control experiment, the impact of GDC-0084 to inhibit AKT phosphorylation in neuronal cultures from α-Syn^A53T^ mice was assessed. AKT phosphorylation at Thr308 and Ser473 was inhibited with GDC-0084, applied at 50 and 100 nM. Moreover, silencing the human A53T α-Syn transgene using shSNCA resulted in lower AKT activation (Fig. 3 C,D).

**Fig. 3.**
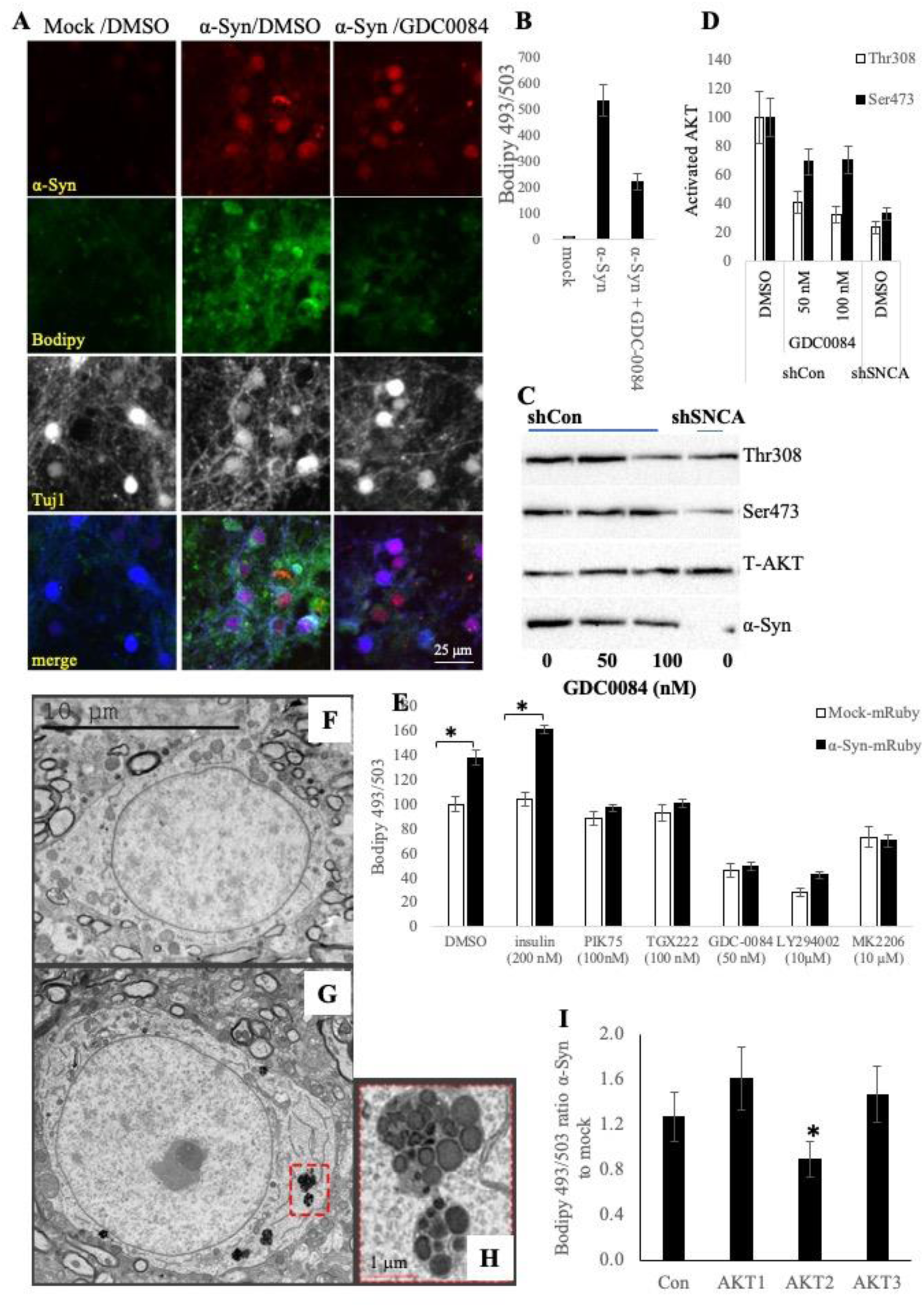
α-Syn-enhanced neuronal lipid droplets (LDs) accumulation is dependent on PI3K/AKT signaling. **(A)** Immunocytochemistry (ICC) of primary cortical neurons from α-Syn^-/-^ mouse brains, infected to express either mock or α- Syn vectors. At 7DIV neurons were treated with GDC-0084 (a PI3K inhibitor, at 50 nM) or solvent (DMSO) for 3 hours, fixed, immunoreacted with anti α-Syn ab (red) and Tuj1 ab (white), and stained for LDs with Bodipy 493/503 (green). **(B)** Bar graph showing quantitative data of mean±SD of n= 12-15 neurons in each treatment. **(C)** GDC-0084 inhibits neuronal PI3K/AKT activation. Primary neurons from α-Syn^A53T^ brains, infected with a scrambled shRNA (shCon) or shSNCA, and treated with GDC-0084 as in (A). AKT phosphorylation detected by Western blotting. **(D)** Bar graph showing mean±SD data (as in C) of n=2 experiments. **(E)** α-Syn -mediated LD-accumulation is sensitive to PI3K/AKT activation/inhibition. HeLa cells infected to express α-Syn-mRuby or a mock- mRuby. 48 hours post infection, cells were treated with insulin for 30 minutes, or with the indicated inhibitors for 3 hours in 10% serum. Control cells treated with DMSO. Detection of LDs with Bodipy 493/503 via FACS. Mean±SD of n>50,000 gated cells (based on the fluorophore). **(F-H)** Representative electron micrographs of coronal brain sections containing a cortical neuron from of an 8 month-old control **(F)** or α-Syn^A53T^ **(G)** mouse brains. Showing accumulation of cytoplasmic lipid droplets **(H). (I)** α-Syn–mediated LDs accumulation is preferably activated by AKT2. HeLa cells co-infected with α-Syn-mRuby or mock-mRuby together with a CRISPR-CAS vector to silence either one of the three AKT isoforms. At 72 hours post infection, lipid droplets were determined following staining of live cells with Bodipy 493/503 via FACS. Graph shows the ratio of Bodipy signal obtained in α-Syn- to mock-expressing cells. Mean±SE of >40,000 gated cells, * P<0.001, one way ANOVA.

The role of α-Syn in promoting LD accumulation and its dependence on the PI3K/AKT pathway was further examined in HeLa cells. Cells expressing either a mock-mRuby or α- Syn-mRuby plasmid were initially analyzed for AKT phosphorylation to ensure that the mRuby fluorophore or the A2 peptide did not interfere with the impact of α-Syn to activate it (Fig. S1). Cells expressing α-Syn or the mock plasmid for 48 hours were subsequently incubated with PI3K inhibitors (e.g., PIK75, TGX221, GDC-0084, LY294002) or an AKT inhibitor (MK2206) for 3 hours. A parallel culture was treated with insulin to activate the pathway. LDs accumulation was determined by FACS in live cells. The results further confirmed that the effects of α-Syn on enhancing the accumulation of LDs are mediated by the PI3K/AKT pathway (Fig. 3E).

To validate the presence of structured neuronal LDs, we examined coronal sections from α-Syn^A53T^ and control WT C57BL/6 mouse brains at eight months of age using transmission electron microscopy (TEM). Characteristic LDs, observed as densely stained clusters within the cytoplasm, were frequently identified in cortical neurons of α-Syn^A53T^ mice, but were absent in the control WT C57BL/6 brain sections (Fig. 3 F-H).

To find out if a specific AKT isoform is preferably involved in α-Syn-mediated LD accumulation we co-expressed in HeLa cells α-Syn or a mock plasmid together with a gRNA directed at either one of the AKT isoforms and the degree of LDs accumulation was determined by FACS. Downregulating either AKT1, AKT2 or AKT3 resulted in ∼ 45% lower mRNA levels. Notably, the impact of α-Syn to enhance LD accumulation over their levels in the control cells was lost in cells that their AKT2 isoform was silenced (Fig. 3I). This suggests that α-Syn preferentially facilitates LD accumulation through AKT2 downstream of PI3K.

### α-Syn-mediated LD accumulation is mTOR and PPARγ-dependent

To investigate the impact of α-Syn on mTOR activation downstream of PI3K/AKT, we analyzed the expression levels of effector proteins for mTORC1 and mTORC2 following pathway activation in primary cortical neurons, dissociated from α-Syn^-/-^ brains, and infected to express α-Syn or a mock vector. The results show a higher degree of phosphorylated S6 (S6P), a downstream mTORC1 effector, in α-Syn- over mock-expressing cells (Fig. 4A,B). Levels of mature SREBP-1 (normalized to actin) and ACLY phosphorylation at Ser455 (normalized to total ACLY) were also elevated with α-Syn expression (Fig. 4A,B). Together with the increases in AKT phosphorylation at Ser473 (Fig. 1), these findings indicate that α-Syn activates both mTOR complexes.

**Fig. 4.**
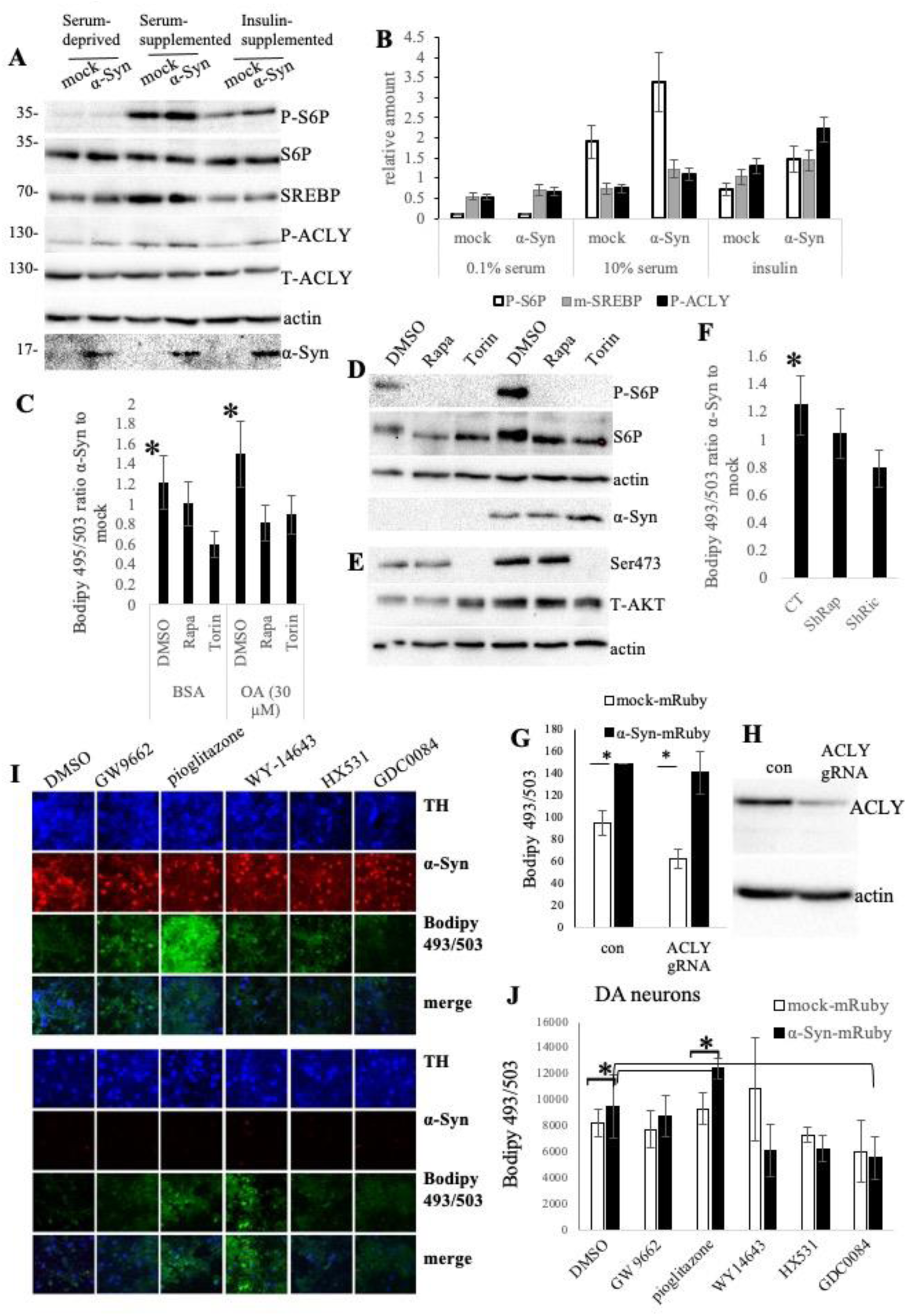
α-Syn-enhanced LDs accumulation is dependent on mTORC1 and enhanced with PPARγ-activation. **(A)** α-Syn activates mTOR complexes. Primary cortical neurons from α-Syn ^-/-^ mouse brains, infected to express α-Syn or mock vectors. At 10 DIV neurons were conditioned in serum-deprived medium (Neurobasal-A) for 16 hours and subsequently transferred to serum-supplemented medium (B27 in Neurobasal-A), or insulin-supplemented Neurobasal-A for additional 20 minutes. Western blot immunoreacted with antibodies against phosphorylated and total S6 (S6P), the mature form of SREBP-1, phosphorylated and total ACLY, actin for loading control, and α-Syn. **(B)** Bar graph as in (A) showing mean ± SD of n=2 repeats. **(C)** Rapamycin (1 μM) or Torin (1 μM) were applied for one hour to HeLa cells infected to express α-Syn-mRuby or mock- mRuby plasmids. The cells were then conditioned in serum-free medium supplemented with either BSA (0.1 mM) or oleic acid (0.03 mM in BSA) for an additional 2 hours in the presence of the inhibitors. LDs were determined by FACS at **48** hours post infection. The ratio of Bodipy signal detected in α-Syn- to mock-infected cells is presented. N>25,000 gated cells. *P < 0.05, one-way ANOVA. **(D, E)** Western blotting performed in parallel to (C) immunoreacted with anti phospho S6P and total S6P. An additional blot reacted with anti Ser473 and total AKT. **(F)** HeLa cells were co-infected to express α-Syn-mRuby or mock-mRuby, along with an shRNA targeting Raptor (shRap) or Rictor (shRic). Cells were serum-starved for 3 hours, followed by serum replenishment for 3 hours, and collected 48 hours post-infection. LDs detected with Bodipy 493/503 via FACS. Data present the mean ± SD from >50,000 gated cells. *P < 0.001. **(G,H)** α-Syn induces LD accumulation independently of ACLY activation. HeLa cells were co-infected with α-Syn-mRuby or mock-mRuby, along with either a gRNA targeting ACLY or a scrambled gRNA (con). Seventy-two hours post-infection, cells were processed for LD detection with Bodipy 493/503. Data present the mean ± SD from >35,000 gated cells. *P < 0.001, t-test. Western blotting immunoreacted with antibodies against ACLY and actin. **(I, J)** Activated PPARγ is required for α-Syn-mediated LD formation. ICC of primary dopaminergic neurons dissociated from α- Syn^-/-^ mouse brains, infected to express α-Syn or a mock-vector. Neurons at 12 DIV were conditioned for 16 hours with GW9662 (50 μM); pioglitazone (50 μM); WY-14643 (100 μM); or HX531 (10 μM). GDC-0084 (70 nM) for 3 hours. Detection of LDs following reaction with anti-Tyrosine Hydroxylase (TH, blue) and anti α-Syn ab (red), and stained with Bodipy 493/503 (green). Graph (J) showing mean ± SD of 40-60 neurons taken at n=3 fields. Signal quantified using QuPath.

To explore the role of mTOR complexes in α-Syn-mediated LD accumulation, HeLa cells expressing α-Syn or a mock plasmid were treated with rapamycin or Torin inhibitors.

Bodipy staining, indicated that both inhibitors reduced α-Syn-mediated LD accumulation (Fig. 4C). Inhibitor efficacy was confirmed by reduced phosphorylation of S6P (mTORC1) and Ser473 (mTORC2) in parallel cultures (Fig. 4D,E). Additionally, downregulating mTORC1 and mTORC2 using shRaptor or shRictor, respectively, also reduced LD accumulation in α-Syn-expressing cells compared to the control cells (Fig. 4F). Efficacy of the shRNAs was validated by decreased S6P and Ser473 phosphorylation (Figs. S2,S3). These results demonstrate that actived mTOR complexes are essential for α- Syn-mediated LD accumulation.

Given ACLY activation upon α-Syn expression (Figs. 4A,B, S4 and S5), and its reported roles in lipogenesis and adipogenesis(*26, 29*), we tested the potential involvement of ACLY in α-Syn -mediated LDs accumulation. However, in experiments in which ACLY expression was significantly downregulated via CRISPR-CAS, we found no effect on α- Syn-mediated LD accumulation (Fig. 4 G,H).

Since α-Syn activates SREBP-1 (Fig. 4A,B), we tested Betulin, an SREBP inhibitor (*33*), in HeLa cells. Betulin effectively blocked α-Syn-induced LD accumulation (Fig. S6).

Among its targets, SREBP-1 specifically activates Peroxisome Proliferator-Activated Receptor (PPAR)γ(*34*). In a previous study, we identified the activation of PPAR isoforms in response to α-Syn expression(*35*). Of the three PPAR isoforms, our results indicated that PPARγ was the most significantly affected by the expression of α-Syn. Given that both mTORC1 and mTORC2 regulate the transcriptional activation of PPARγ-target genes (*28, 29*) which are involved in glucose and lipid regulation(*26, 29*), we sought to investigate whether activated PPARγ is necessary for α-Syn-induced LD accumulation.

Primary dopaminergic neurons from α-Syn^-/-^ mouse brains were infected to express either α-Syn or a mock vector. Neurons were treated with a PPARγ antagonist (GW9662); PPARγ agonist (pioglitazone); PPARα agonist (WY14643); RXR antagonist (HX531); or GDC-0084 (PI3K inhibitor). Cells were then fixed, immunoreacted with anti-Tyrosine hydroxylase (TH), a marker for dopaminergic neurons and anti α-Syn antibodies, and stained with Bodipy 493/503. The results confirmed that α-Syn enhances LD accumulation in TH-positive dopaminergic neurons (Fig. 4I,J). Notably, pioglitazone enhanced the Bodipy 493/503 signal in α-Syn-expressing neurons, while GW9662 inhibited α-Syn-induced LD accumulation. WY14643 enhanced the signal only in the mock vector-expressing cells. Similar results, showing a higher degree of Bodipy 493/503 signal in α-Syn expressing cells treated with pioglitazone were obtained in cortical neurons from α-Syn^A53T^ brains via ICC (Fig. S7) and also in HeLa cells via FACS (Fig. S8). These results indicate that α-Syn facilitates LD accumulation downstream of PI3K/AKT/mTOR and PPARγ activation.

### α-Syn stabilizes/facilitates palmitoylated Ras localization at the plasma membrane

The sk-Mel2 cell line is derived from a malignant melanoma of human origin and expresses detectable levels of endogenous α-Syn. Attempting to silence α-Syn in these cells, we found no effect for α-Syn expression on PI3K/AKT activation (Fig. S9). This result is in contrast to the effect of silencing α-Syn in neuronal cells, that reduced the degree of AKT activation (Fig. 1E,F). Sk-Mel2 cells harbor a N-Ras Q61R mutation within its GTP-binding region, producing a constitutively activated NRas. The absence of an effect for silencing α-Syn expression on PI3K/AKT activation in these cells has prompt us to hypothesize that α-Syn may associate with Ras to activate PI3K. Considering previous reports for α-Syn associations with FAs and with membranes(*3*), we further hypothesized that α-Syn may associate with palmitoylated Ras proteins at the plasma membrane.

To examine α-Syn’s association with Ras proteins, we co-expressed in HeLa cells α-Syn together with HRas-GFP or NRas-GFP. Control cells were similarly co-infected but with a mock plasmid. α-Syn-positive cells showed enhanced Ras localization to the plasma membrane (Fig. 5A,B). Mutating palmitoylation sites in HRas (Cys181,184) or NRas (Cys181) abolished this effect. Fractionation of α-Syn-/- cortical neurons expressing α- Syn or mock vectors confirmed increased membranal Ras with α-Syn expression (Fig. 5C,D).

**Fig. 5.**
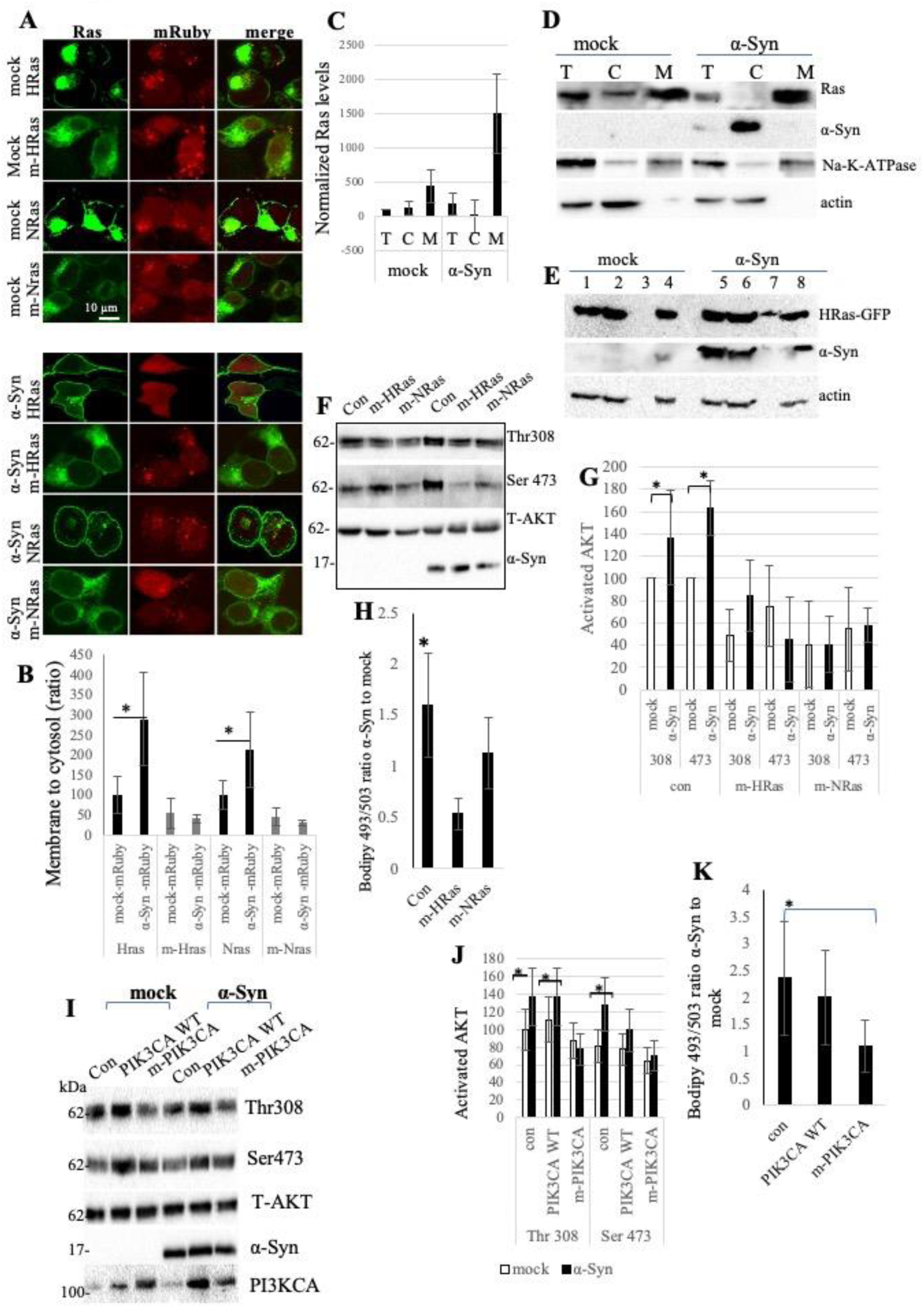
α-Syn stabilizes/facilitates palmitoylated Ras at the plasma membrane to activate PI3K p110α. **(A)** Hela cells co-infected with mock-mRuby or α-Syn- mRuby together either with HRas-GFP, NRas-GFP, mutant HRas (m-HRas, C181S and C184S), or m-NRas (C181S). Forty-eight hours post infection, cells were fixed and process for direct visualization of the flourophores. **(B)** Graph showing quantitative data of the Ras-GFP signals presented as a ratio between the membranal (measured as 2.5 μm in width relative to the outer ring surrounding the cell) and cytosolic fraction in each cell. Quantitative results obtained with QuPath and are mean±SD of n=12-15 cells in each group. *, p> 0.05. **(C, D)** Western blot analysis of cytosolic (C) or membrane (M) fractions and total cell lysates (T) from cultured primary neurons infected to express α-Syn or a mock plasmid. Immunoblotted with antibodies against Ras, Na-K-ATPase and actin. Representative blot out of n=2. Graph shows mean ± SD of n=2 independent experiments. **(E)** Representative immunoblot of S-palmitoylation assay for HRas- GFP. HeLa cells, expressing a mock vector or α-Syn were serum-starved over- night followed by 30 minutes activation in serum-replenished medium. Cells were collected 72 hours post infection and snap-froze. Palmitoylated HRas-GFP captured by CAPTUREome resin is detected in cleaved bound fraction of α-Syn expressing cells (cBF, lanes 3 and 7). Input fraction (IF, lanes 1 and 4) represents total cell extract, and the flowthrough fraction represents the cleaved unbound protein (cUF, lanes 2 and 6). The negative control samples, treated with acyl- preservation reagent and subjected to incubation with CAPTUREome resin is preserved bound fraction (pBF, lanes 4 and 8). **(F-H)** Palmitoylated Ras is required for the impact of α-Syn to enhance AKT activation and LD-accumulation. HeLa cells co-infected to express α-Syn-mRuby or mock-mRuby together with m-HRas or m-NRas. Cells were serum starved for 16 hours and then serum-replenished for one hour. Western blot showing AKT activation levels in lysed cells 48 hours post infection **(F).** A representative blot. Bar graph showing quantitative analysis of n=3 blots **(G).** LDs determined by FACS, presenting the ratio of Bodipy signal in α-Syn to mock-infected cells. Mean ±SD of >35,000 gated cells, *, P< 0.05, one way ANOVA **(H). (I-J)** Hela cells co-infected with a mock-mRuby or α-Syn- mRuby together with WT PIK3CA or mutant PIK3CA (m-PIK3CA), carrying a dual T208D and K227A mutation at the RBD. AKT activation determined 48 hours post infection by Western blotting (a representative blot out of n=4, mean± SD, *, P<0.05, one way ANOVA. **(K)** HeLa cells as in (I) analyzed for LDs. Presenting the ratio of Bodipy signal in α-Syn to mock-infected cells. Mean±SD of >50,000 gated cells. Mean± SD, *, P<0.05, one way ANOVA.

As an additional indication for the impact of α-Syn on membrane localization of Ras, we determined its impact on palmitoylated HRas-GFP protein, migrating at 48 kDa. Using an assay detecting S-palmitoylation, we found a higher degree of cleaved-bound HRas-GFP in HeLa cells co-expressing α-Syn and HRas-GFP than in the control cells (Fig. 5E).

We examined whether Ras palmitoylation is essential for α-Syn-induced AKT phosphorylation (Fig. 5F,G) and LD accumulation (Fig. 5H) in HeLa cells co-expressing α-Syn or mock vectors, together with mutant HRas or NRas. The results confirmed that α- Syn requires palmitoylated Ras to promote both effects.

We next analyzed the impact of the palmitoylation inhibitor 2-bromopalmitate (2-BP) and the APT-1/APT-2 inhibitors ML348 and ML349, which facilitate Ras depalmitoylation(*36*), on α-Syn -mediated LD accumulation. 2-BP effectively blocked LD accumulation in α-Syn-expressing cells, while ML348 and ML349 had no effect (Fig. S10). Additionally, 2-BP suppressed α-Syn-induced increases in PIP3 and PI4,5P2 (Fig. S11, S12), supporting a role for protein palmitoylation in α-Syn’s activation of PI3K/AKT and LD formation.

### RBD in PI3KCA is required for α-Syn-mediated LD accumulation

Activated Ras proteins bind directly to the RBD of p110α to activate its lipid kinase activity(*17, 37*). To find out whether α-Syn -dependent PI3K/AKT activation requires Ras-bound p110α, we introduced T208D and K227A point mutations in the RBD of PIK3CA to interfere with Ras binding without affecting PIK3CA catalytic activity(*17*). Cells were infected to co-express α-Syn together with WT- or mutant-PIK3CA, or a control (con) vector. Sister cultures were similarly co-infected but with a mock vector for α-Syn. We found no detectable effect for over expressing WT-PIK3CA, however, mutant PIK3CA (m-PIK3CA) prevented α-Syn from activating AKT at Thr 308 and Ser 473 positions (Fig. 5 I,J). In complement, m-PIK3CA expression abolished the enhancing effect of α-Syn on LD accumulation. That is, the Bodipy 493/503 signal detected in α-Syn over mock-infected cells was lower in cells expressing m-PIK3CA (Fig. 5K).

### PI3K activation is associated with PSer129 α-Syn levels

To assess PI3K/AKT pathway involvement in α-Syn pathophysiology, we analyzed PSer129 α-Syn levels in cortical neurons from α-Syn^A53T^ mice. Neurons were serum-starved overnight, then either maintained in starvation medium (control) or treated with serum or insulin to activate PI3K/AKT signaling. Parallel cultures were treated with the PI3K inhibitor GDC-0084 (50 or 100 nM) or DMSO in serum-replenished media. AKT phosphorylation at Thr308 and Ser473 was analyzed (Fig. 6A-D), and PSer129 α-Syn levels were quantified by Lipid-ELISA as a percentage of total α-Syn(*38*) (Fig. 6E-F). Notably, PSer129 α-Syn levels correlated positively with AKT phosphorylation.

**Fig. 6.**
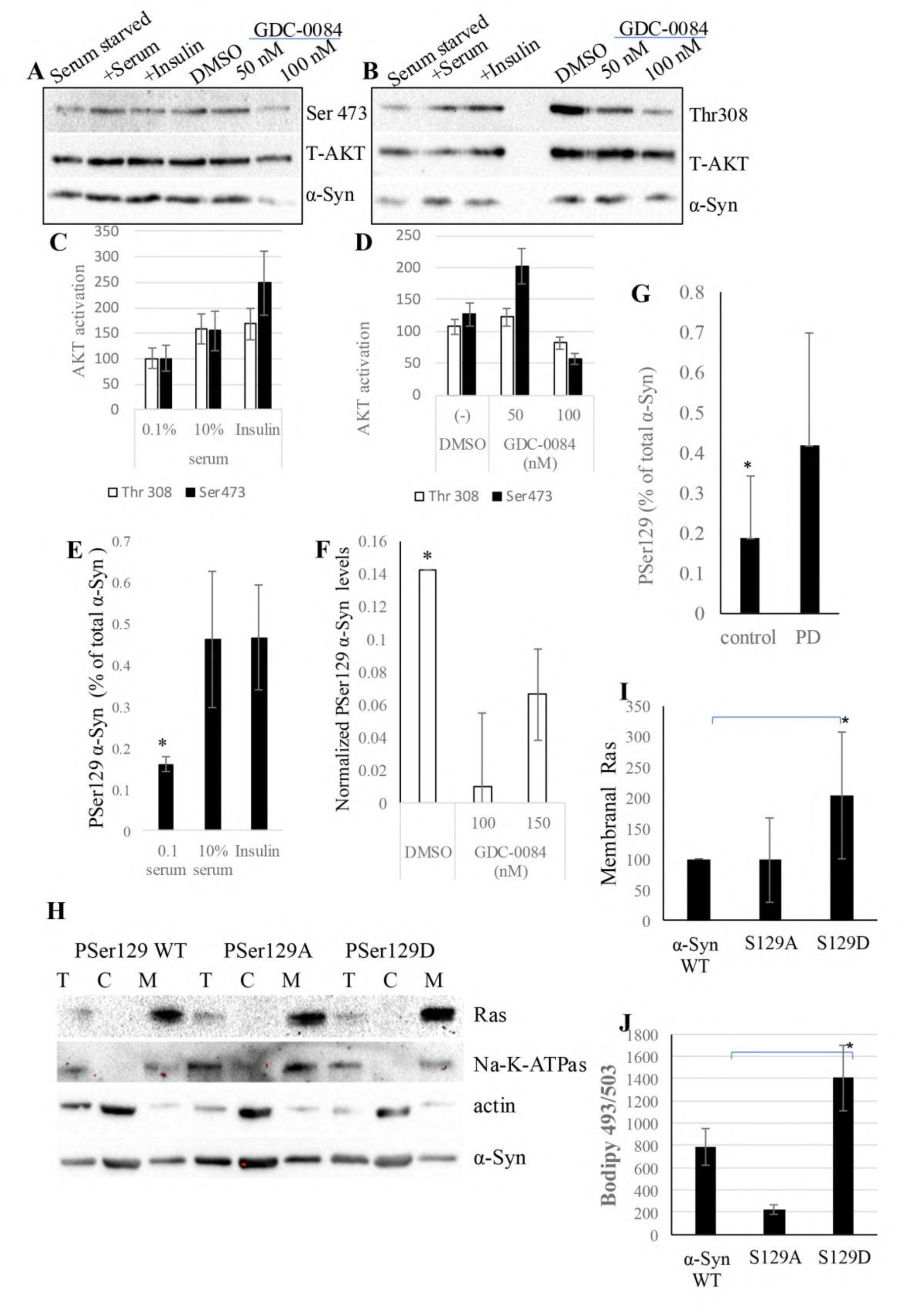
PSer129 α-Syn levels correlate with PI3K/AKT Activation. **(A, B)** Primary cortical neurons dissociated from α-Syn^A53T^ mouse brains. At 10 DIV neurons were serum-starved overnight in Neurobasal-A medium without B27 supplement, followed by 15-minute activation with either B27 serum or Neurobasal-A containing insulin .). Alternatively, neurons were treated with the GDC-0084 inhibitor (at 50 and 100 nM) or DMSO (solvent) for 2 hours. Western blotting immunoreacted with antibodies against Thr308-AKT, Ser473-AKT, total AKT, and α-Syn. A representative blot is shown (out of n=2). **(C,D)** Quantitative analysis (for blots in A and B) showing mean ± SD of n=2 independent experiments. **(E, F)** PSer129 α-Syn detection by Lipid ELISA. The graph shows mean ± SD of n=3 independent experiments. PSer129 α-Syn levels are presented as percent of total α-Syn detected in the same sample. **(G)** Levels of PSer129 α- Syn normalized to total α-Syn determined by Lipid ELISA in the soluble fraction of lysed caudate from PD and control human brains. Graph showing mean±SD of n=12 brains in each group. *, P>0.01, ttest. **(H, I)** HeLa cells infected to express WT α-Syn, Ser129Ala or Ser129Asp and fractionated to collect cytosolic (C) membrane (M) and total cell lysates (T). Immunoblotted with antibodies against Ras, Na-K-ATPase and actin. Bar graph shows Mean±SD on n=3 repeat. **(J)** HeLa cells as in (H) analyzed for LDs.

We next measured PSer129 α-Syn in the soluble, cytosolic fraction of human brain tissue (caudate) from PD and controls, without dementia, and expressed it as a percentage of total α-Syn levels. This ratio was ∼0.18% and 0.42% in control and PD brains, respectively (n=12 brains each, Fig. 6G). Notably, these values are in the range of PSer129 α-Syn levels following PI3K/AKT activation in the primary neurons (Fig. 6 E,F).

To assess the involvement of PSer129 α-Syn in PI3K activation, we expressed in HeLa cells either WT α-Syn, a Ser129Ala mutation that hinders phosphorylation, or a Ser129Asp mutation, considered chemically similar to phospho-serine. We find a significantly higher degree of membrane associated Ras proteins following cell fractionation (Fig. 6H,I) and a higher degree of LDs (Fig. 6J) in cells expressing the phosphomimetic Ser129Asp mutation.

### Investigating the effects of GDC0084 on α-Syn-related pathologies in α-Syn^A53T^ mice

Progress in the development of PI3K inhibitors in clinical use and the availability of GDC-0084, a PI3K inhibitor that was shown to cross BBB (*39*), has prompted us to investigate the impact of this drug on α-Syn- pathologies in-vivo, in α-Syn^A53T^ and control α-Syn^-/-^ mice. Mice at 4-5 months of age, including males and females, were randomly divided in four groups, to receive GDC-0084 at 20 mg/kg/day or the vehicle for 28 consecutive days, as previously described (*40*). Drug’s activity in the brain was determined based on levels of phosphorylated AKT at Thr308 and Ser473 (Fig. 7 A,B). Showing a lower degree of Thr308 AKT in the drug- treated groups. In addition, levels of Ser 473 AKT were found lower in drug-treated α-Syn^-/-^ brains but not in the α-Syn^A53T^ brains. The presence of the drug in brains of treated α-Syn^A53T^ as well as control mice was confirmed by mass spectrometry (Fig. 7 C).

**Fig 7.**
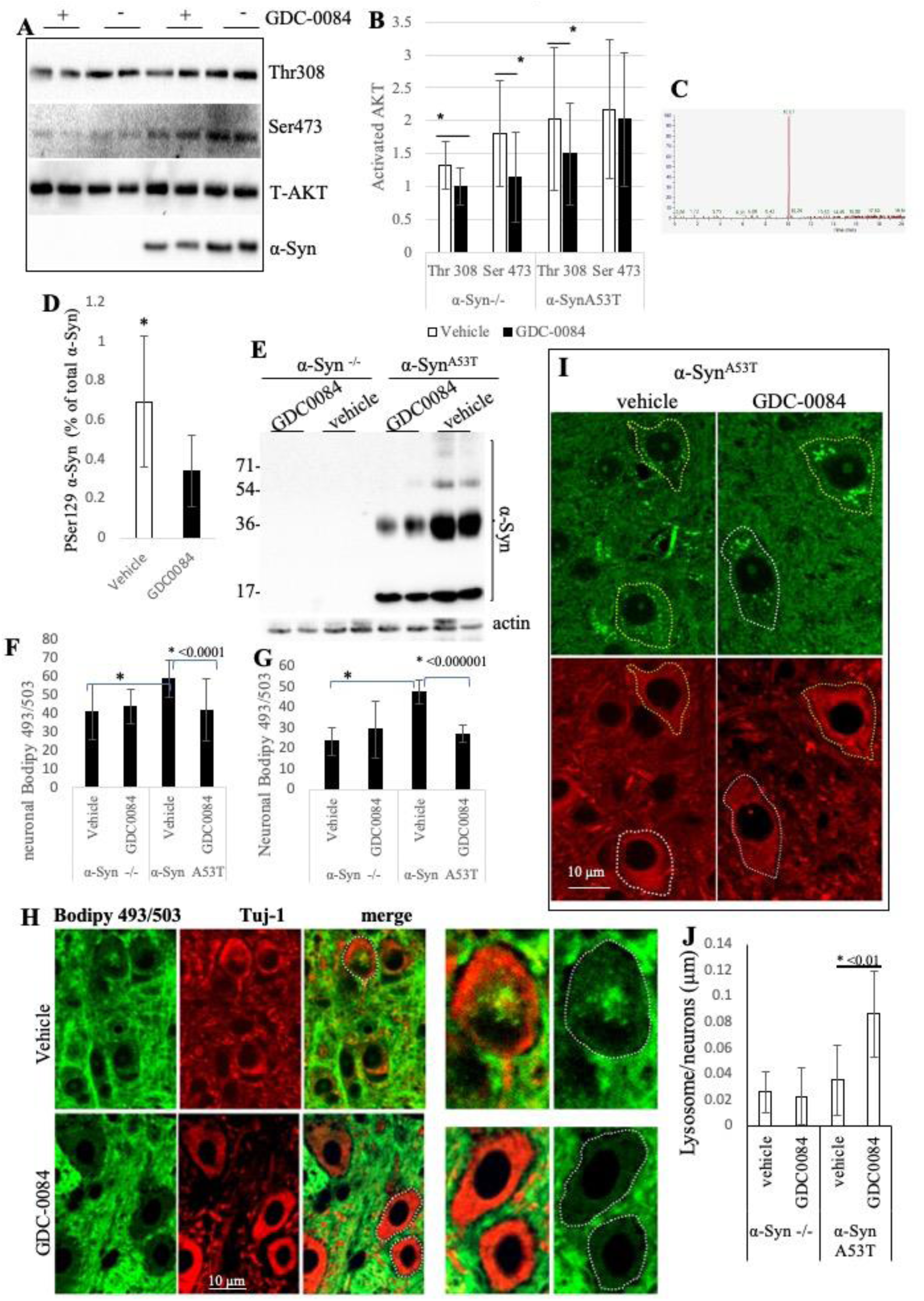
Beneficial effects for GDC-0084 treatment in α-Syn^A53T^ mice. **(A)** α- Syn^A53T^ or control α-Syn^-/-^ mice at 4-5 months of age, including males and females, were randomly divided in four groups, to receive GDC-0084 at 20 mg/kg/day or the vehicle for 28 consecutive days. Western blot analysis of proteins extracted from one brain hemisphere (including left and right hemispheres) of α- Syn^A53T^ or control α-Syn^-/-^ brains. A representative blot out of n=4. **(B)** Bar graph showing mean±SD on n=6 different brains in each group. *, P<0.05, One way ANOVA. **(C)** Mass spectrometry data demonstrating the detection of GDC-0084 in a mouse brain treated with the drug. **(D)** levels of PSer129 α-Syn detected in the soluble fraction of α-Syn^A53T^ mouse brains (vehicle- or drug-treated) determined by Lipid ELISA. Mean±SD of n=6 α-Syn^A53T^ brains in each group, analyses include either left or right hemispheres, females and males (with equal presentation). Data presented as percent PSer of total α-Syn. * p< 0.01, ttest. **(E)** Detection of α-Syn oligomers following incubation with Lipidex. Protein sample from brain homogenates (soluble fraction, 50 μg) analyzed by Western blotting and immunoreacted with anti α-Syn ab. A representative blot out of n=3. **(F-H)** Paraffin-embedded brain sections co-stained with Bodipy 493/503 and Tuj1 antibody. Graphs showing quantitative neuronal Bodipy 493/503 signal detected in the cortex (F) or brain-stem (G). (H) Bodipy 493/503 signal (green) detected within neurons stained with Tuj1 ab (red) from α-Syn^A53T^ mouse brains, treated with vehicle (upper panel) or with GDC-0084 (lower panel). **(I)** IHC of paraffin-embedded brain sections co-stained with anti LAMP1 (green) and Tuj1 (red) antibodies. **(J)** Bar graph showing size of LAMP-1 positive clusters as a ratio of the size of neurons in GDC-0084 or vehicle treated mice. Mean±SD of n>25 cells in each group. *, P<0.01, ttest.

**Fig. 8.**
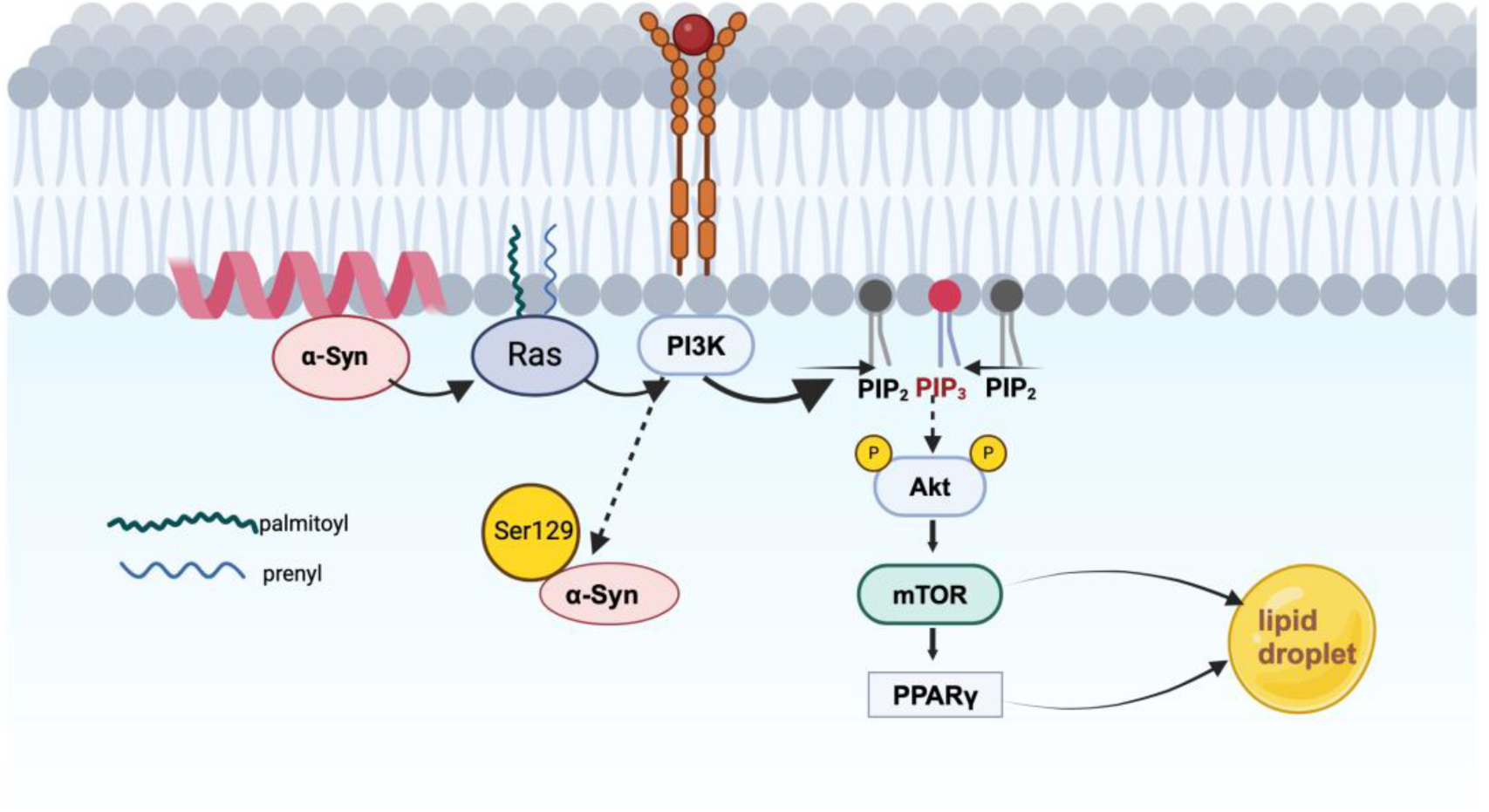
graphical abstract. The PI3K/AKT pathway is activated in PD. α-Syn facilitates the membrane localization of palmitoylated Ras proteins, which are effector proteins of PI3K. The activated PI3K/AKT/mTOR pathway results in the accumulation of LDs, as part of disease pathology.

PSer129 α-Syn levels were found to be lower in the soluble fraction of GDC-0084-treated α-Syn^A53T^ brains (Fig. 7 D). Furthermore, treatment with GDC-0084 also lowered the levels of soluble α-Syn oligomers detected in the soluble fraction of drug-treated α- Syn^A53T^ brains (Fig. 7 E).

To examine the impact of GDC-0084 on accumulation of neuronal LD, mediated by α- Syn, we co-stained paraffin-embedded brain sections, with Bodipy 493/503 and Tuj1 antibody, a neuronal marker, focusing on two brain regions, brain-stem and cortex. Consistent with findings from cultured neurons (Figs. 3 and 4, above), a higher neuronal Bodipy signal was observed in α-Syn^A53T^ compared with α-Syn^-/-^ brains, treated with the solvent only. Notably, following GDC-0084 treatment, neuronal Bodipy signal was reduced in the α-Syn^A53T^ mouse brains, resulting in comparable Bodipy signals between the two genotypes and across the tested brain regions (Fig. 7 F-H). To analyze the appearance and distributions of lysosomes within neuronal cells, we co-stained paraffin-embedded brain sections with anti Lamp1 and Tuj1 antibodies (Fig. 7 I,J). Neuronal lysosomes which are immunoreactive with Lamp1 were found to be clustered close to the perinuclear region in brain sections of α-Syn^A53T^ treated with GDC-0084. However, the lysosomes in neurons from vehicle-treated α-Syn^A53T^ were distributed throughout the cells (Fig. 7 I,J). No differences were detected in lysosomal dispersal in drug- or vehicle-treated control α-Syn^-/-^ brain sections. Clustering of lysosomes has been previously reported in PD and disease models(*41–43*) and lysosomal clusters are associated with induced autophagy (*44, 45*) as well as clearance of pathogenic α-Syn species (*46, 47*). Together, GDC-0084 treatment in α-Syn^A53T^ mice normalized the hyperactivated PI3K/AKT signaling, lowered the accumulation of LD, decreased α-Syn species associated with he pathogenesis of PD to levels comparable to control mouse brains, and promoted lysosomal clustering.

## Discussion

This study identifies a role for α-Syn in activating the PI3K/AKT pathway by promoting Ras localization at the plasma membrane. Data obtained in young (and apparently healthy) α-Syn^A53T^ mouse brains and from dissociated cortical neurons, silenced for their endogenous α-Syn expression, support the conclusion that the role of α- Syn in activating PI3K/AKT is part of its normal physiological function in the healthy brain. Moreover, evidence for pathway hyper activation, detected in PD brains, underscores relevance to disease mechanisms. Focusing on LDs we report that α-Syn enhances the accumulation of LDs downstream of PI3K/AKT/mTOR and PPARγ activation. This effect was consistently observed across multiple systems, including cell lines, mouse cortical and dopaminergic neurons. Treatment with GDC-0084 showed beneficial effects in the α-Syn^A53T^ mouse model. Including, management of PSer129 α- Syn levels, α-Syn oligomerization, LD accumulation and lysosomal dispersion. These findings collectively highlight a regulatory role for α-Syn in the interplay between PI3K/AKT activation and the neurodegenerative processes characteristic of PD, opening new avenues for therapeutic developments.

### α-Syn, PI3K/AKT/mTOR signaling, and its dual role in neurodegeneration and melanoma

Pathological α-Syn has been shown to activate mTORC1, leading to impaired autophagy in cellular and animal models of PD (*21*) and rapamycin, an mTORC1 inhibitor, has demonstrated neuroprotective effects by mitigating dopaminergic neuron loss (*21*). Additionally, mTORC1 inhibition via a CCR5 antagonist has been reported to exert protective effects by modulating microglial-to-neuronal signaling, enhancing autophagic clearance, and reducing neurotoxicity in neurodegeneration (*48*). These findings highlight mTORC1, a regulator of autophagy, as a potential therapeutic target for PD and other neurodegenerative diseases. Our data indicating PI3K activation by α-Syn suggest that, beyond mTORC1, additional upstream components of this signaling cascade, including PI3K itself, AKT, and mTORC2, may contribute to α-Syn-related pathologies. Supporting evidence may include alterations in PIP levels observed in PD and related models(*49*), excessive axonal growth and arborization associated with the disease (*50*), mitochondrial dysfunction and oxidative stress, synaptic dysfunction, and disruptions in dopamine signaling (*20, 51, 52*). Thus, understanding the contribution of these upstream components is crucial for elucidating disease mechanisms and identifying therapeutic targets.

Furthermore, hyperactivation of the PI3K/AKT pathway in PD brains and α-Syn-expressing disease models aligns with evidence of a mechanistic link between PD and melanoma. Epidemiological studies indicate an increased risk of melanoma in PD patients (*53*). Notably, the PI3K/AKT/mTOR pathway plays a significant role in melanoma progression (*54*) and α-Syn is expressed at elevated levels in melanoma and other cancers (*55, 56*). Suppression of α-Syn expression in melanoma reduces tumor growth (*57*), whereas α-Syn oligomers and phosphorylated Ser129 α-Syn have been associated with increased tumor cell proliferation (*55*).

### α-Syn as a regulator of membrane localization of Ras

The results indicate that the localization of HRas and NRas at the plasma membrane is facilitated and/or stabilized by the expression of α-Syn. Given that Ras signaling relies on its GDP-GTP exchange for effector protein activation, proper Ras localization at the plasma membrane is critical for activating class I and class II PI3Ks, which contain Ras-binding domains(*58–61*). The precise mechanism by which α-Syn facilitates Ras localization at the plasma membrane remains unclear. One potential mechanism may involves Ras palmitoylation, a process that serves as a membrane anchor for HRas, NRas, and the KRas4A isoform. Notably, a recent study have linked α-Syn to palmitate turnover in disease models, suggesting its possible involvement in this process (*62, 63*). Another plausible mechanism relates to α-Syn’s role in vesicular trafficking. α- Syn may act to facilitate the vesicular transport of palmitoylated Ras from the Golgi to the plasma membrane. Supporting this hypothesis, Rab27b, a member of the Rab family of small GTPases, has been shown to regulate NRas palmitoylation and its trafficking to the plasma membrane(*64*). Considering Rab27b’s role as a regulator of α-Syn toxicity (*65*), the findings support the need for further research to better understand the impact of α-Syn on protein palmitoylation, in particular, Ras palmitoylation, and the relationship to disease.

### PSer 129 α-Syn and PI3K/AKT activation in neuronal activity

Phosphorylation of α-Syn at Ser129 is a post-translational modification linked to α-Syn toxicity in PD(*66*). It has also been described as a dynamic and physiological post translational modification driven by physiological neuronal activity(*67*). Unlike the non-phosphorylated α-Syn, PSer129 α-Syn is detected only in a subset of brain regions (*68, 69*), and this phosphorylation enhances synaptic targeting and protein-protein interactions (*68, 69*). Despite efforts to establish a pathological role for PSer129 α-Syn, no clear link between phosphorylation, aggregation, and toxicity has been found (*66*). The rapid changes in PSer129 α-Syn levels in response to PI3K/AKT activation (occurring within minutes of activation) and the correlation between PSer129 α-Syn levels and the degree of PI3K/AKT activation, observed in human brains and neurons from α-Syn^A53T^ mouse brains, support a role for PSer129 α-Syn in cellular mechanisms regulated by the PI3K/AKT pathway. Furthermore, these findings indicate that PSer129 α-Syn may contribute to PI3K/AKT-dependent mechanisms in the healthy brain and in neurodegeneration.

### Study limitations and interpretation

While our findings provide strong evidence for α-Syn-induced activation of the PI3K/AKT pathway and its role in lipid droplet accumulation, several limitations should be considered. First, although we establish a mechanistic link between α-Syn, PI3K/AKT activation, and lipid metabolism, the broader implications for neuronal survival and disease progression remain to be fully elucidated. Additionally, our study focuses on the PI3K/AKT pathway and does not address the potential involvement of the MAPK/ERK pathway downstream of Ras. Translating these findings into clinical applications will require rigorous validations.

## Materials and Methods

**Human brains** frozen brain tissue containing the caudate from advanced PD (Braak stage 5-6) and age-matched control brains, were obtained from by the Multiple Sclerosis Society Tissue Bank, funded by the Multiple Sclerosis Society of Great Britain and Northern Ireland, registered charity 207495. The approval for the use of human tissue material was obtained from the Peer Review Panel of the Parkinson’s UK Brain Bank.

### Mice

The human PrP-A53T α-Syn tg mouse line (α-Syn^A53T^)(*70*) was purchased from Jackson Laboratory (Bar Harbor, ME, USA) as hemizygous; cross-bred with α-Syn^-/-^ C57BL/6JOlaHsd mice (Harlan Laboratories, Jerusalem, Israel(*71*)) to remove endogenous mouse α-Syn; and then bred to achieve homozygosity of the human A53T α- Syn transgene. α-Syn^-/-^ C57BL/6JOlaHsd (*71*) and WT C57BL/6JRccHsd genotypes are used for control mouse lines. All animal welfare and experimental protocols were approved by the Committee for the Ethics of Animal Experiments of the Hebrew University of Jerusalem NIH approval # OPRR-A01-5011 (Permit number: MD-16- 14826-3).

### Cell cultures

HEK 293T and HeLa cell lines were maintained in Dulbecco’s modified eagle’s medium (DMEM) supplemented with 10% FBS; 2% L-glutamine; 1% penicillin/streptomycin, sodium-pyruvate and non-essential amino acids (Biological Industries, Beit-Haemek, Israel). SK-mel2 cells experiments were performed between weeks 2-4 from thawing a frozen aliquot to maintain optimal endogenous α-Syn expression. Cells were maintained in minimum essential medium (MEM; Sigma-Aldrich, Rehovot, Israel) supplemented with 10% FBS; 1% L-glutamine, penicillin/streptomycin and sodium-pyruvate. HepG_2_ cells were maintained in minimum essential medium (EMEM) supplemented with 8% FBS, 1% L-glutamine, penicillin/streptomycin and sodium-pyruvate. Cultures were maintained at 37°C in a 95% air/ 5% CO2 humidified incubator.

In experiments involving serum-starvation cells were starved overnight in 0.1% FBS, followed by the indicated treatment added to cells for the time indicated in the presence of serum.

## Reagents

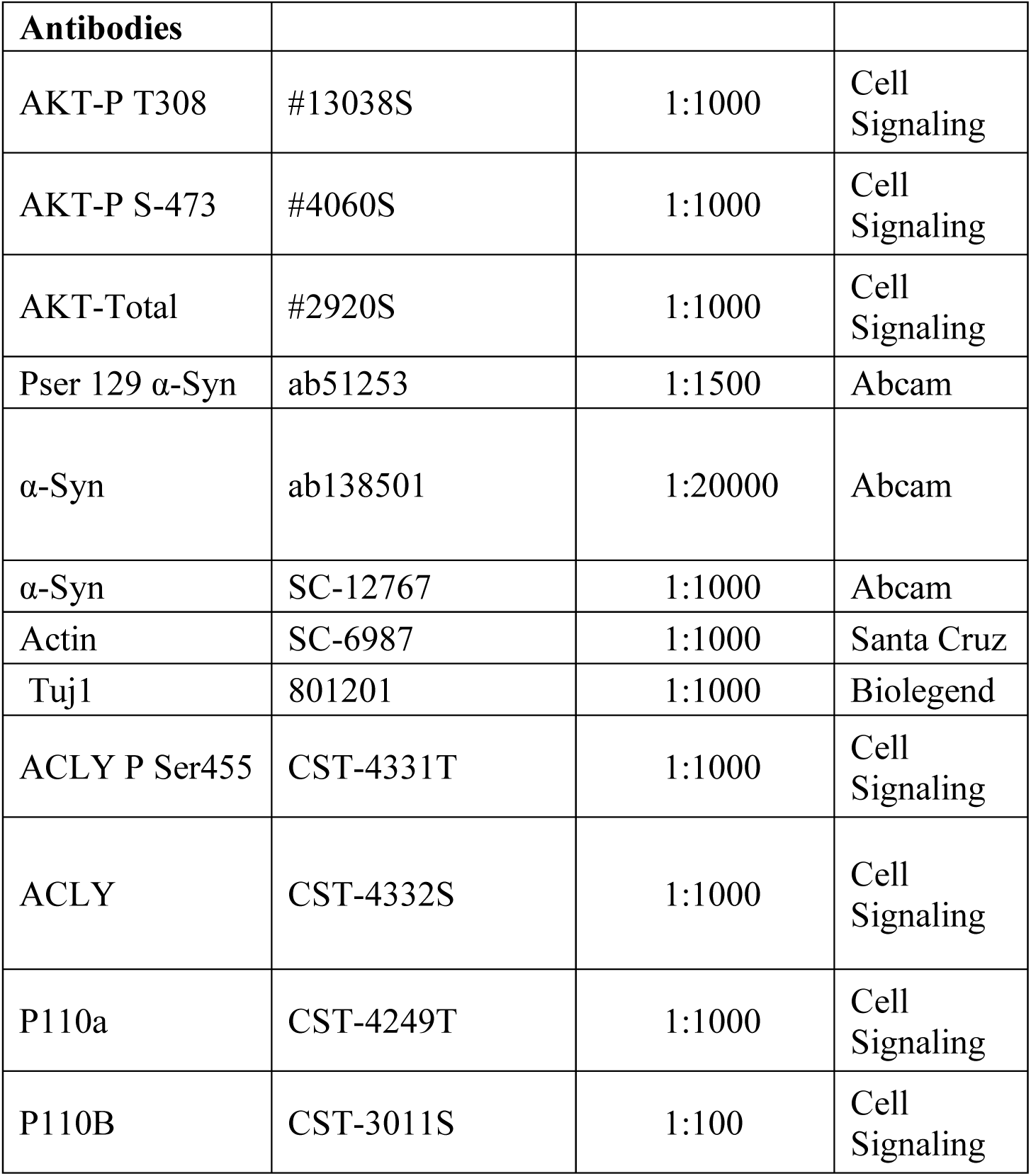
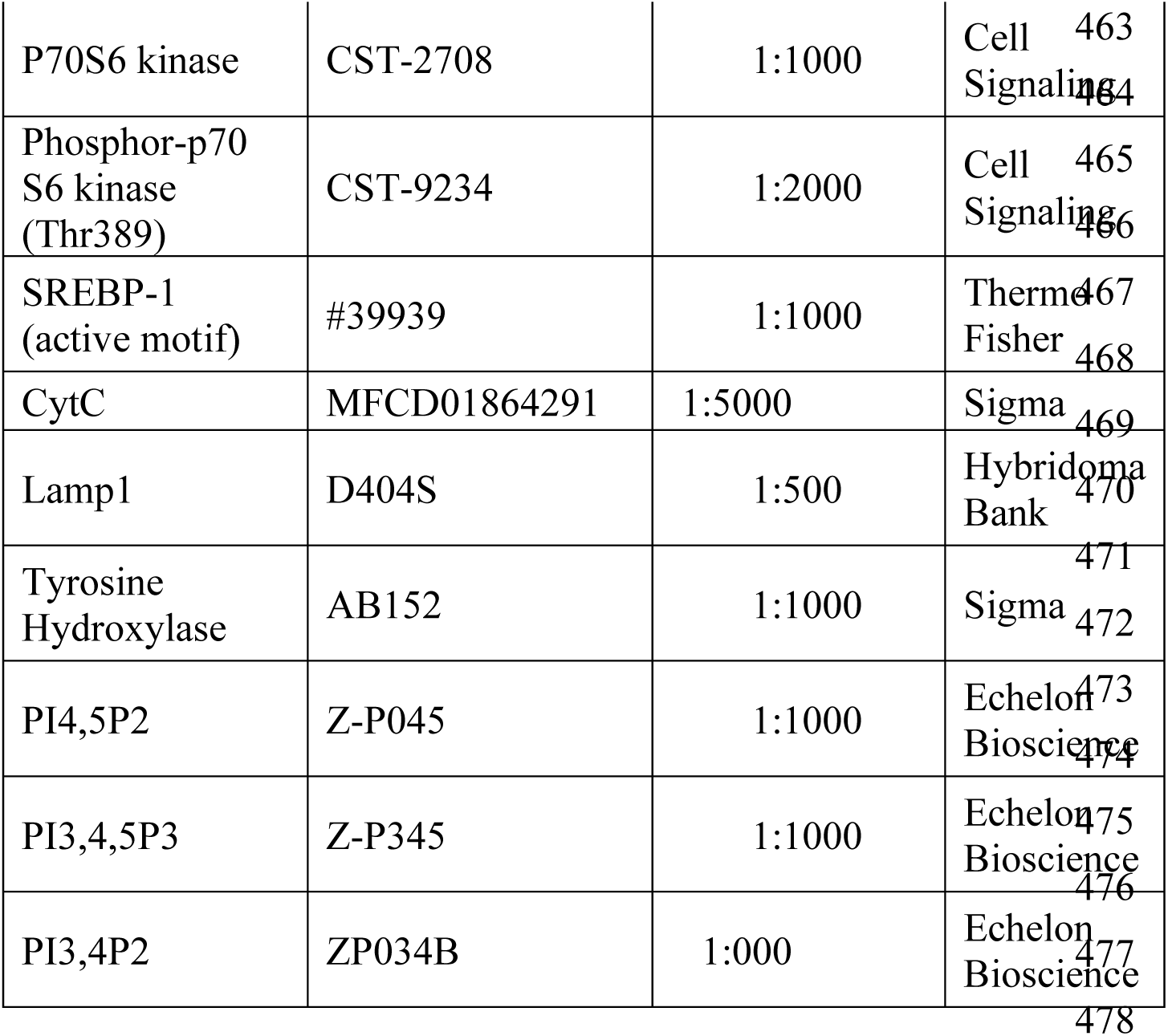

## Chemaicals

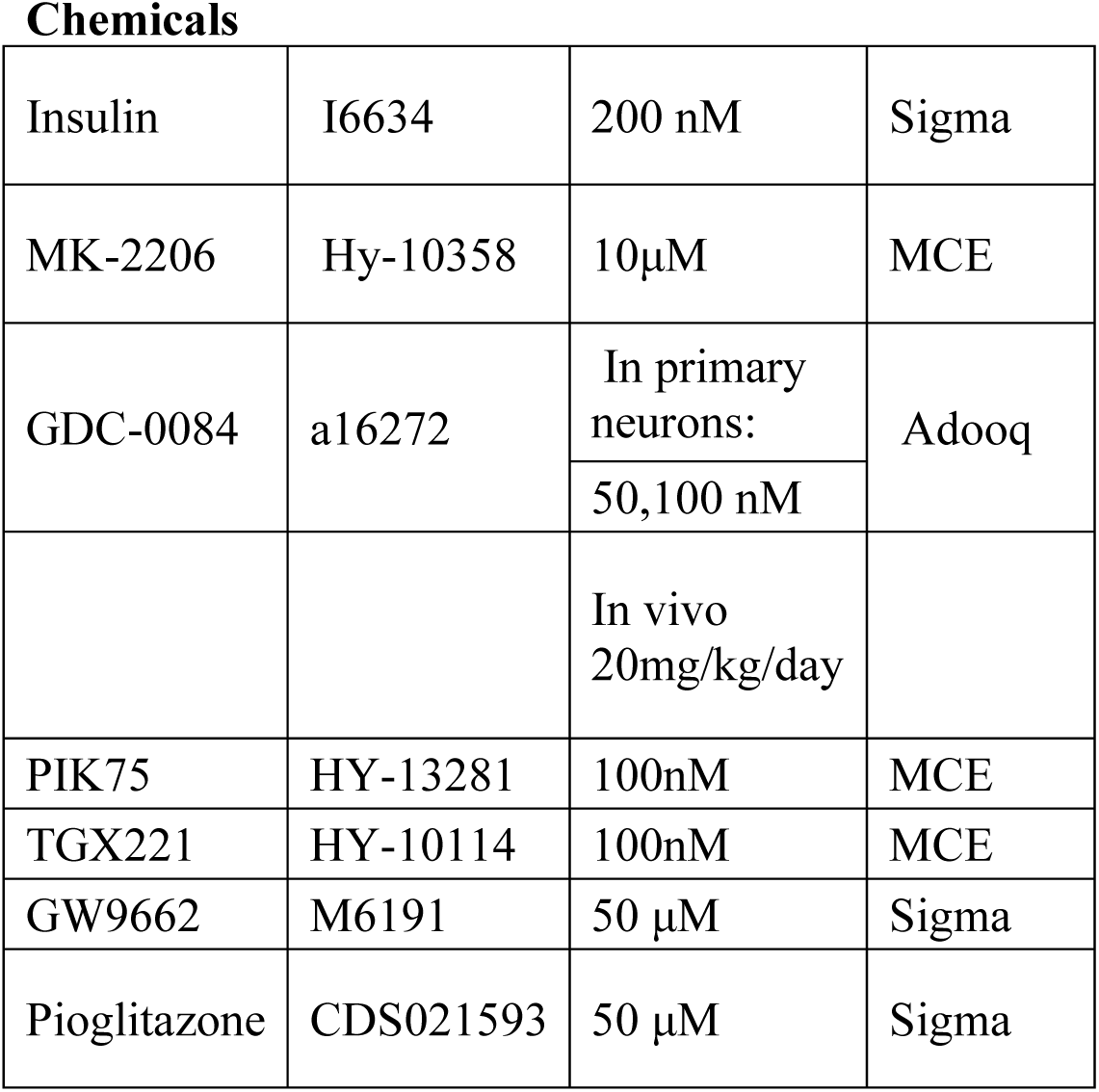
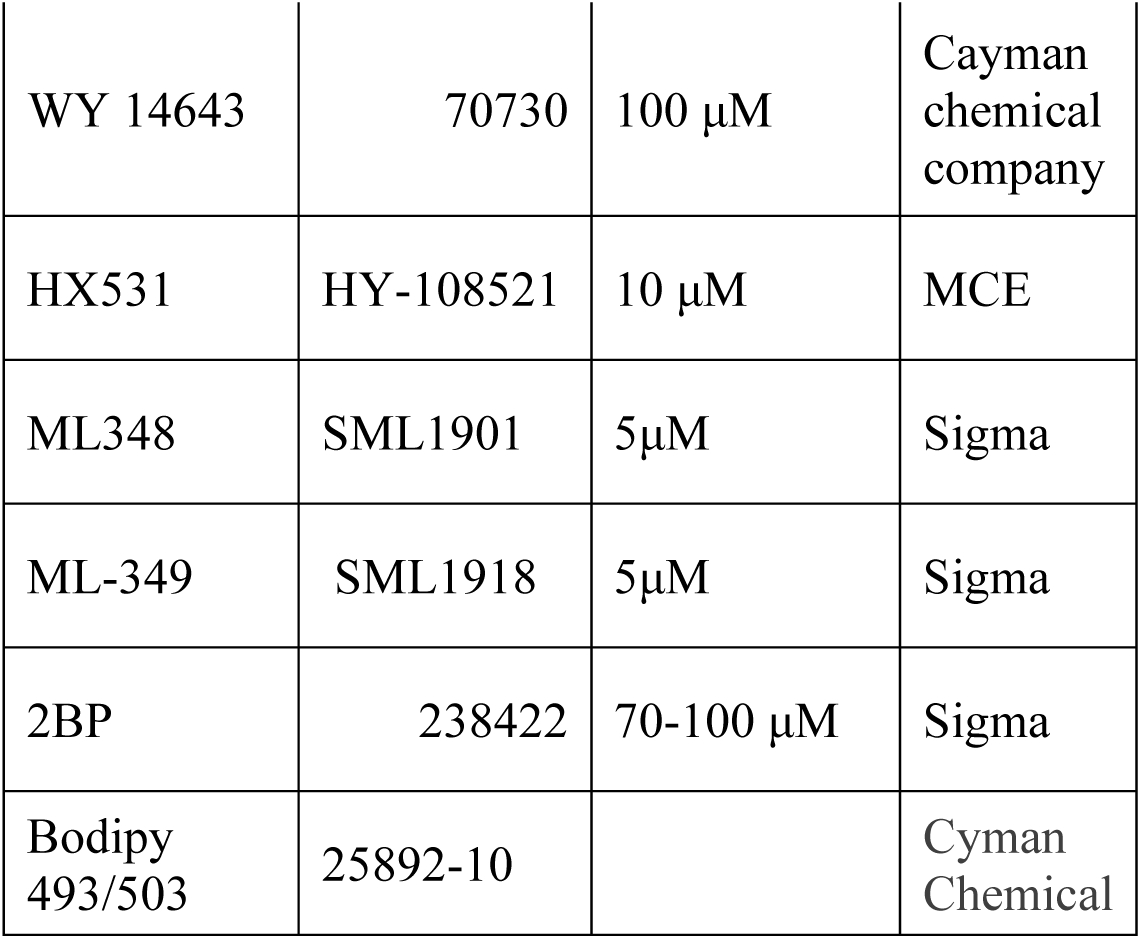

Chemicals
|  |  |  |  |
| --- | --- | --- | --- |
| Insulin | I6634 | 200 nM | Sigma |
| MK-2206 | Hy-10358 | 10µM | MCE |
| GDC-0084 | a16272 | In primary neurons: | Adooq |
|  |  | 50,100 nM |  |
|  |  | In vivo<br>20mg/kg/day |  |
| PIK75 | HY-13281 | 100nM | MCE |
| TGX221 | HY-10114 | 100nM | MCE |
| GW9662 | M6191 | 50 µM | Sigma |
| Pioglitazone | CDS021593 | 50 µM | Sigma |
| WY 14643 | 70730 | 100 μM | Cayman chemical company |
| HX531 | HY-108521 | 10 μM | MCE |
| ML348 | SML1901 | 5μM | Sigma |
| ML-349 | SML1918 | 5μM | Sigma |
| 2BP | 238422 | 70-100 μM | Sigma |
| Bodipy 493/503 | 25892-10 |  | Cyman Chemical |

## Plasmids

### Mutant PIK3CA

WT PIK3CA (donated by Bert Vogelstein, Addgene 16643) was cut using Acl-I and Bgl-II and the DNA fragment was replaced with a sequence containing the RBD mutations that was synthesized at SynBio Technologies (NJ, USA). The DNA fragment was inserted in to introduce T208D (replacing ACT to GAT at position 622-624) and K227A (AAA to GCA at position 678-681) mutations. Both plasmids, WT PI3KCA and m-PIK3CA were cloned into FUGW backbone using Gibson Assembly Master Mix NEB-E2611S.

### α-Syn m-Ruby

A DNA fragment containing mRuby3-P2A (839 bp) was PCR amplified using primers that introduce Age1 restriction sites at both ends (donated by from Tobias Meyer, Addgene #85146). The fragment was ligated upstream of SNCA sequence in FUGE plasmid to generate mRuby-P2A-α-Syn.

### Mutagenesis in Ras palmitoylation sites

WT HRas-GFP (*72, 73*) cloned into FUGW. Mutations (C181S and C184S) introduced by Gibson Assembly^®^ Master Mix (New England Biolabs) using PCR primers forward: 5’gactctagaggatccccgggtaccgATGGTGAGCAAGGGCGAGGAGC 3’; and revers: 5’ cgataagcttgatatcgaattTCAGGAGAGCACACACTTGCTGCTCATGCTGCCGG 3’

WT NRas-GFP^115^ cloned into FUGW. Mutation (C181S) introduced as above using the following primers: forward: 5’ gactctagaggatccccgggtaccgATGGTGAGCAAGGGCGAGGAGC 3’; and reverse: 5’ cgataagcttgatatcgaattTTACATCACCACACATGGCAATCCCATACTACCCTGAGTCC C 3’.

Ras-iRFP670 was cloned by removing GFP and inserting the iRFP amplified from pNLS-iRFP670 (donated by Vladislav Verkhusha, addgene #45466).

### CRISPR /Cas9 cloning

a guide RNA (gRNA) sequences for CRISPR /Cas9 was designed at the official CRISPR website (http://crispr.mit.edu) and based on previous reports. The insert gRNA oligonucleotide for the genes of target and the complementary oligonucleotides are annealed and cloned into lenti crispr V2 vector (Addgene #52961). For virus production, HEK293T cells are transfected with a control, scrambled gRNA or a specific gRNA. Lentiviruses are purified, titered, concentrated and stored in aliquots. The infection efficacy is generally ranging between 75-90%. The following gRNAs were used: PIK3CA: 5’ACAGCCACACACTACATCAG; PIK3CB: 5’ ATGCATATCTGATTTTACGA; ACLY:5’ACCAGCTGATCAAACGTCGTGG;

AKT1 gRNA (BRDN0001146935); AKT2 gRNA (BRDN0001148949); and AKT3 gRNA (BRDN0001146868) donated by John Doench & David Root, addgene # 75500, addgene # 77506, and addgene #76217.

### shRNA

Costume-ready Mission shSNCA were from Sigma-Aldrich (TRCN0000272292), shRictor, (Addgene #1853, donated by David Sabatini); shRaptor (Addgene #1857, donated by David Sabatini).

### Primary cultures from mouse brains

Cortical cultures were prepared from cortices, dissected from a day old (P1) C57BL/6JRccHsd or C57BL/6JOlaHsd mouse brains, as described previously ^51^. Briefly, cells (∼50,000) were plated onto coverslips, pre-coated with 12.5 μg/ml poly-D-lysine (Sigma-Aldrich) in a 12-well dish. Cortical neurons were maintained in Neurobasal-A medium (Gibco, Thermo Fisher Scientific, Petah Tikva, Israel) and supplemented with 2% B-27 (Gibco, Thermo Fisher Scientific); 1% L-glutamine; 0.5% penicillin/streptomycin. To eliminate glia cells, 1μM cytosine β-D- arabinofuranoside (Ara-C; Sigma-Aldrich) was added to the culture at 1-2 DIV. Culture medium was partially (25-50%) replaced every 4 days. Cultures were maintained at 37°C in a 5% CO2 humidified incubator. Mesencephalic cultures were prepared from brains of E13.5 mouse embryos as described(*74*) and plated onto poly-D-lysine coated coverslips. Cells were infected immediately at 1DIV.

**Transmission electron microscopy** was performed as described previously (*50*). Briefly, 100 μm coronal brain sections were obtained using a vibratome (Leica Biosystems, IL, USA). Brain sections were fixed in Karnovsky’s fixative solution. Sections were washed four times with 0.1M sodium cacodylate buffer (pH 7.3) and incubated for 1 h in 1% osmium tetroxide, 1.5% potassium ferricyanide in sodium cacodylate. Sections were then washed; dehydrated with graded series of ethanol solutions (30, 50, 70, 80, 90,95, 100%); followed by propyleneoxide. Brain sections were infiltrated with series of epoxy resin, (25, 50, 75, 100%) and polymerized to generate blocks that were sectioned by an ultramicrotome (Ultracut E, Riechert-Jung, Ontario,Canada). 80 nm sections were stained with uranyl acetate and lead citrate. Sections were observed using a Jeol JEM 1400 Plus Transmission Electron Microscope and pictures were taken using a Gatan Orius CCD camera.

### α-Syn and Pser129 α-Syn detection by Lipid ELISA

The method as described (*38*) was slightly modified. Lipid mix (phosphatidylinositol (PI), phosphatidylserine (PS), phosphatidylethanolamine (PE), and GM-1 ganglioside (Sigma, Rehovot, Israel) at a 1:1:1:1 ratio) were dissolved in methanol. Total α-Syn detection using MJFR1 anti α-Syn ab (Abcam ab138501) at 1:20000 dilution, followed by a one-hour incubation with a secondary anti-rabbit antibody (Jackson ImmunoResearch 711-035-152). Pser129 α-Syn detection, using phospho S129 antibody (Abcam ab51253) 1:1500 dilution, and a secondary anti-mouse antibody (Jackson ImmunoResearch 715-035-150).

Sample preparation: soluble fraction prepared by homogenizing cells or tissue in homogenization buffer (*75*) supplemented with phosphatase inhibitors (5 mM NaPPi, 50 mM NaF), and a protease and phosphatase inhibitor cocktail (Tivan Biotech). Samples were applied to a 96-well Lipid ELISA plate in triplicate at 10-40 ng protein/well after dilution with 1% BSA (fatty-acid free) in PBS (MgCl_2_ and CaCl_2_ free). Membrane fraction(*75*) was resuspended in a lysis buffer containing 50 mM Tris-HCl, pH 8.5, 1% Triton, 1 mM EGTA, 1 mM EDTA, 150 mM NaCl, 5 mM NaPPi, 50 mM NaF, and a protease and phosphatase inhibitor cocktail (Tivan Biotech). Samples of membrane proteins were applied to the Lipid ELISA plate in triplicate at 0.25-1 μg/well after dilution with 1% BSA in PBS (as above).

PSer129 α-Syn levels in human brains are detected in the soluble fraction at the picogram scale (per microgram sample protein) and are normalized to total α-Syn levels detected at the nanogram scale (per microgram sample protein).

### Immunocytochemistry (ICC)

ICC was performed as described previously(*50*). Briefly, cell lines or primary neuronal cultures were fixed in cold 2% paraformaldehyde for 20 min, washed in PBS, and permeabilized with 0.5% saponin in blocking solution (1% BSA in PBS (w/v)) for 30 min at room temperature. Cells were incubated with the primary antibodies (Tuj1:1000, TH 1:1500, Lamp-1 1:100) overnight at 4 °C. Cells were then washed (PBS; 10 min ×3) and incubated with a host-suitable secondary ab, washed again, and mounted in Vectashield mounting medium (Vector Laboratory, Burlingame, CA USA). Images were captured using Nikon’s A1R+ confocal microscope magnification x60, Nikon Spinning Disk Microscope magnification x40.

### Immunohistochemistry (IHC)

IHC was performed as described previously(*50*). Briefly, paraffin-embedded coronal mouse brain sections (6 μm) were processed for immunostaining as previously described(*50*). Antigen retrieval for Lamp-1 and Tuj1 was performed by heating the slides in 10 mM citrate buffer (pH 6) to 95°C for 20 minutes in a pressure cooker. Antigen retrieval for Tuj1 and Bodipy 493/503 staining was performed by incubating the slides with 10 mM citrate buffer (pH 6) at 37°C for 15 minutes. Slides were then blocked with blocking solution (5% normal horse serum in 1% BSA with 0.1% Tween-20) for 1 hour at room temperature, and incubated with primary antibodies diluted in the blocking solution overnight (Tuj1 1:1000, Lamp1 1:100). Slides were washed and incubated with the host-matching secondary antibodies. Images acquired using Nikon Spinning Disk Microscope at x 60 magnification. Quantification of neuronal Lamp1 -positive lysosome: Signal quantified using QuPath. The area of a neurons was identified based on Tuj1 signal (Cy3). Using the option of pixel classifier, a constant threshold was subtracted and LAMP1 signal (FITC) that appears in structures larger than 2 microns in diameter within the neurons was identified. The quantitative data shows the area ratios of lysosome out of the neuron area. N= 5 images from each brain (n > 4 cells in each image). Quantification of neuronal Bodipy 493/503 signal. Images were captured using Cytation 5 (Agilent, BioTek) at x40 magnification. The area of a neuron was identified and annotated based on Tuj1 signal (Cy3) and FITC signal within the area was quantified. N= 8 images from each brain (n > 7 cells in each image).

### Detection of PIP by FACS

Analysis was performed as previously described(*50*). Briefly, cells were fixed in 2% paraformaldehyde at 4°C and permeabilized in 0.2% saponin in 1% BSA (w/v) for 15 minutes at 4°C. For PIPs detection, cells were then incubated with anti-PIP antibodies (Echelon Biosciences), including anti PI(4,5)P2 (Z-A045 1:800), anti PI(3,4,5)P3 (Z- P345b; 1:300) or anti PI(3,4)P2 (Z-P034b 1:500), for 90 minutes with gentle rolling, then washed and probed with the respective secondary antibody, Goat anti-mouse IgG H&L Alexa Fluor 488 (ab150113) or Goat anti mouse IgM Alexa Fluor 488 (ab150121) for 30 minutes at room temperature. Gating for α-Syn expressing cells based on the expression of Ruby P2A α-Syn plasmid. The CytoFLEX LX1 Analyzer was used to analyze the samples.

**Lipid droplets** in cultured cell lines: Cells were stained with Bodipy 493/503 as described previously(*76*). Cells were washed three times with PBS, followed by incubation with 2 μM Bodipy for 15 minutes at 37°C. After three washes with PBS, the cells were trypsinized and filtered with a cell strainer for FACS analysis.

Bodipy staining for ICC or IHC: slides were processed for immunodetection and following the incubation with the secondary antibody, slides were incubated with 1 μg/ml Bodipy staining solution in PBS for one hour at 37°C.

#### S-Palmitoylation Assay

Thiol side-chain protection assay was performed using CAPTUREome™ S-Palmitoylated Protein Kit (Badrilla) in accordance with manufacturer’s instructions.

#### LCMS metabolomics

Analysis was performed as follows: Dionex Ultimate 300 high-performance liquid chromatography (UPLC) system coupled to an Orbitrap Q-Exactive *plus* Mass spectrometer (Thermo Fisher Scientific). Electrospray ionization in the HESI source in a positive mode. The UPLC setup included a C18 column (Kinetex 2.6μm). Five µL of the tissue extracts were injected and the compound was separated with a mobile phase gradient of 15 min, starting at 1% A (H2O:MeOH and 0.1% formic acid) and 99% B (MeCN +0.1% formic acid). The gradient increased linearly from 1% to 75% mobile phase B. This composition was then maintained at 75% Mobile Phase B for the next 5 minutes, from 10 to 15 minutes. The flow rate and column temperature were maintained at 0.2 mL/min and 35 °C, respectively, for a total run time of 30 min. GDC- 0084 was detected with a mass accuracy of less then 5 ppm. Data acquisition was performed using Thermo Xcalibur. Data processing and absolute quantitation were conducted using Trace Finder by Thermo Fischer.

## Statistical Analysis

Appropriate statistical analysis using either Student’s T-test or one-way ANOVA with Dunnett’s multiple comparisons test were used to determine statistical difference, and P values < 0.05 were considered as statistically significant (Prism 7, GraphPad).

## Acknowledgments

We thank Drs. Eduard Berenshtein and Zakhariya Manevitch from the core facility at Hebrew university for professional assistance with electron microscopy and spinning disk microscopy.

## Author contributions

All authors had full access to the data in the study and accepted responsibility to submit it for publication. S.AE. and R.S conceived the study and designed experiments.

S. AE., J.C-L., E.H., D.H., L.N., and M.S performed cellular and/or biochemical studies. O.S., A.E. performed Mass Spect analyses. S. AE., J.C-L., D.H., L.N., performed in vivo study and analyzed data. S. AE., and R.S, writing – review & editing. R.S., supervision and funding acquisition.

## Competing interests

A provisional patent: "PI3K INHIBITORS FOR TREATMENT OF α-SYN-RELATED PATHOLOGIES" was filed Feb 2025. RS.

During the preparation of this work the authors used ChatGPT in order to edit language of the manuscript. After using this tool/service, the authors reviewed and edited the content as needed and take full responsibility for the content of the published article.

## Data and materials availability

All data are available in the main text or the supplementary materials.

---

**S1.**
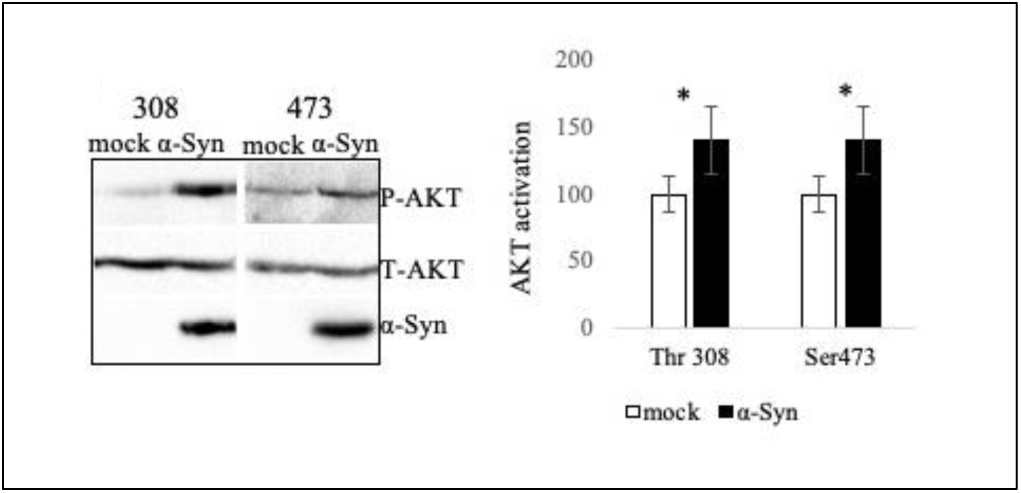
α-Syn-mRuby activates AKT in HeLa cells. Hela cells infected to express a mock- mRuby or α-Syn-mRuby plasmids. Forty-eight hours post infections, cells were lysed and processed for the detection of phospho-AKT by Western blotting, using antibodies against Thr308, Ser473 or total AKT abs (Cell Signaling, Ornat Israel). Graph showing mean±SD of n>4 experiments. *, P<0.05, ttest.

**S2.**
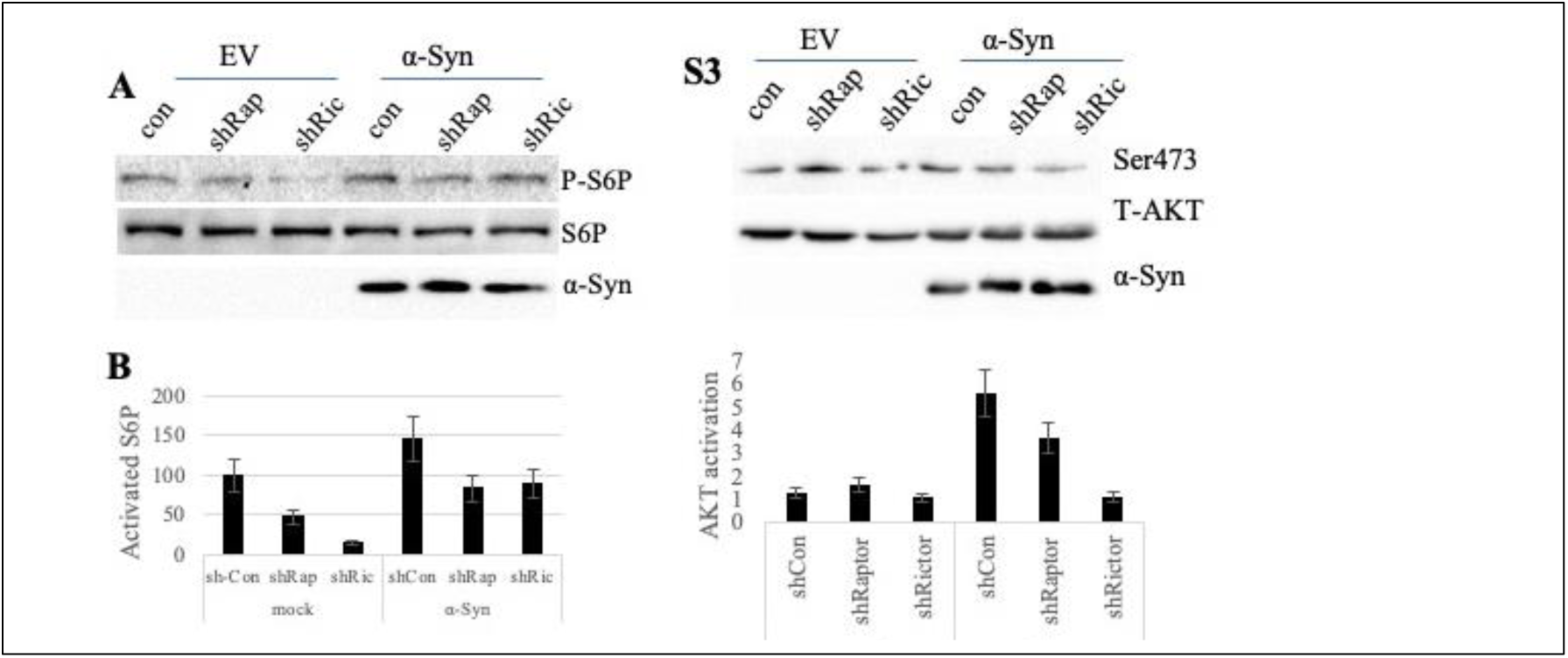
Expression of shRictor/shRaptor in HeLa cells. Cells were co-infected to express α- Syn-mRuby or mock-mRuby, along with a shRNA targeting Raptor (shRap) or Rictor (shRic). Cells were serum-starved for 3 hours, followed by serum replenishment for 3 hours, and collected 48 hours post-infection. Western blot immunoreacted with anti phospho S6P, total S6P and α-Syn antibodies. N=2 repeats. **S3.** Cells as in S2. Western blot immunoreacted with the indicated abs. N=2 repeats.

**S4.**
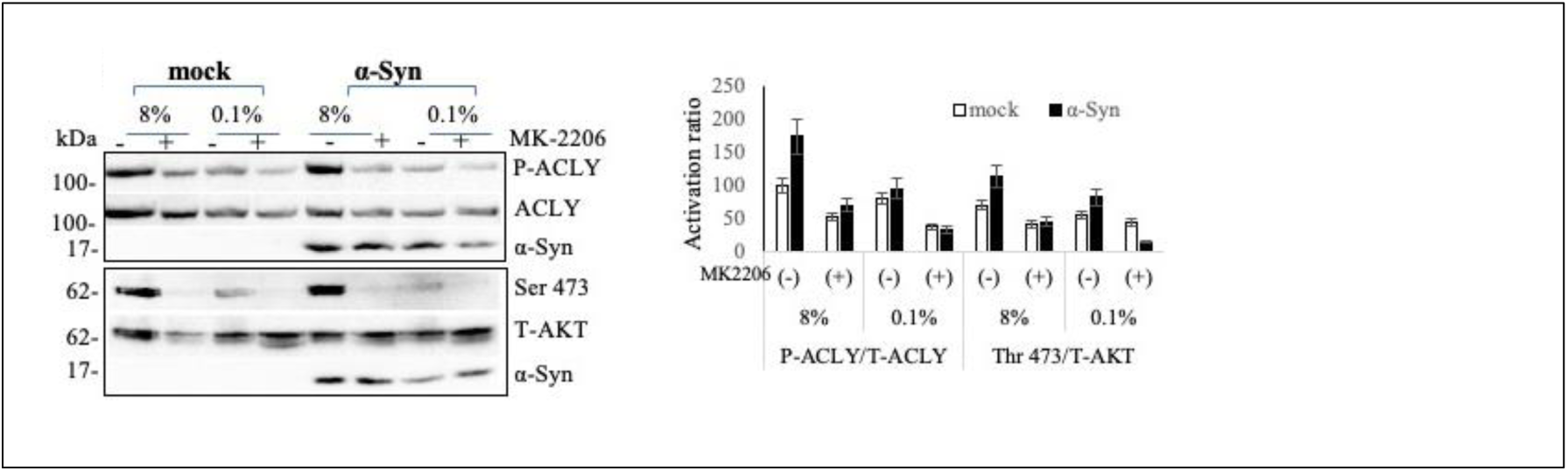
ACLY activation with α-Syn expression requires activated AKT. HeLa cells infected to express α-Syn or a mock vector. 36 hours post infection cells were conditioned for 16 hours in serum supplemeted (10%) or serum-starved (0.1%) media together with MK2206 inhibitor for AKT (10μM) or DMSO (solvent). Representative blot out of n=2 Bar graph showing mean ± SD of n=2.

**S5.**
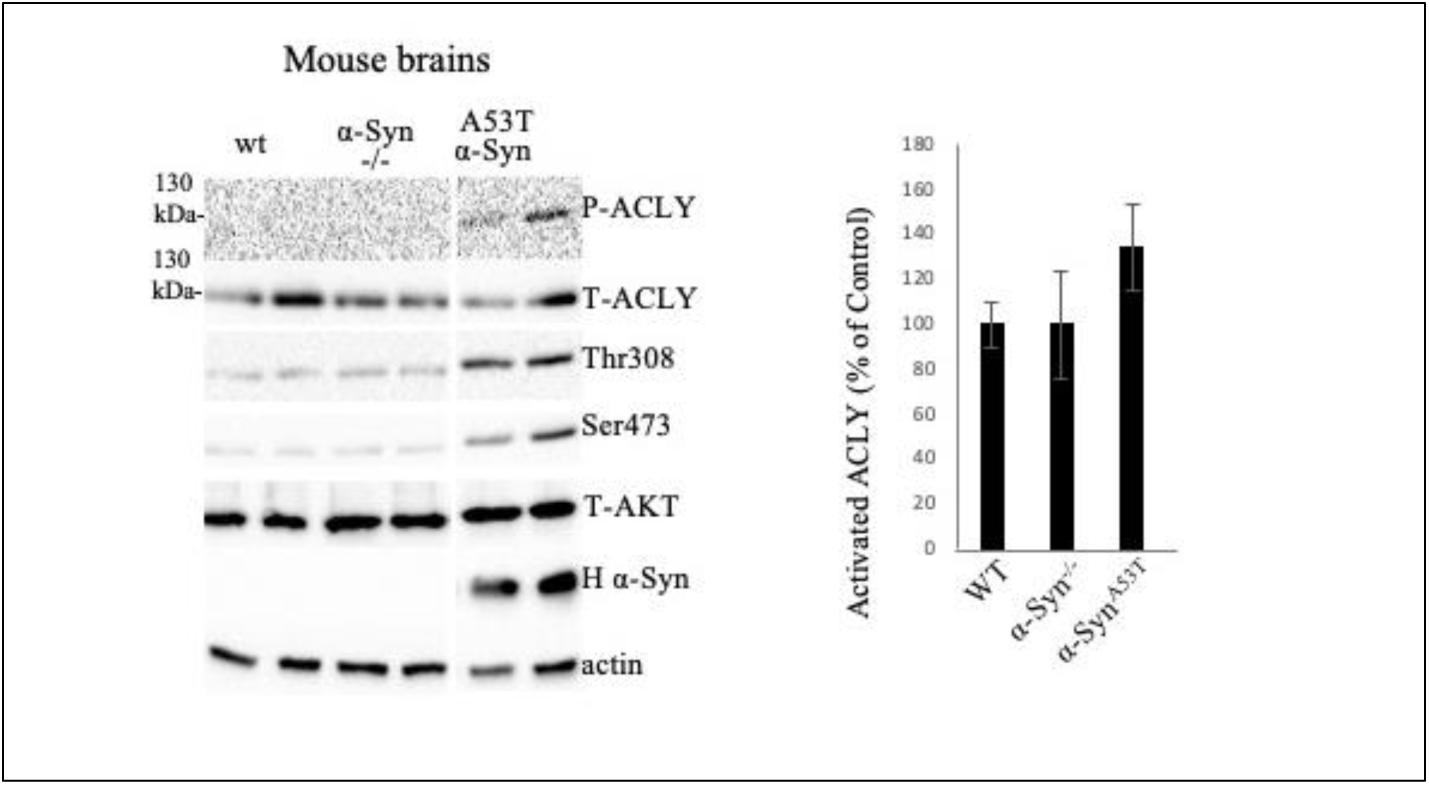
ACLY is activated in brains of α-Syn^A53T^ mice. Whole brain homogenates of mice at 4-6 months old, analysed by Western blotting. Activated ACLY presented as a ratio of phospho-ACLY at Ser 455 to total ACLY.

**S6.**
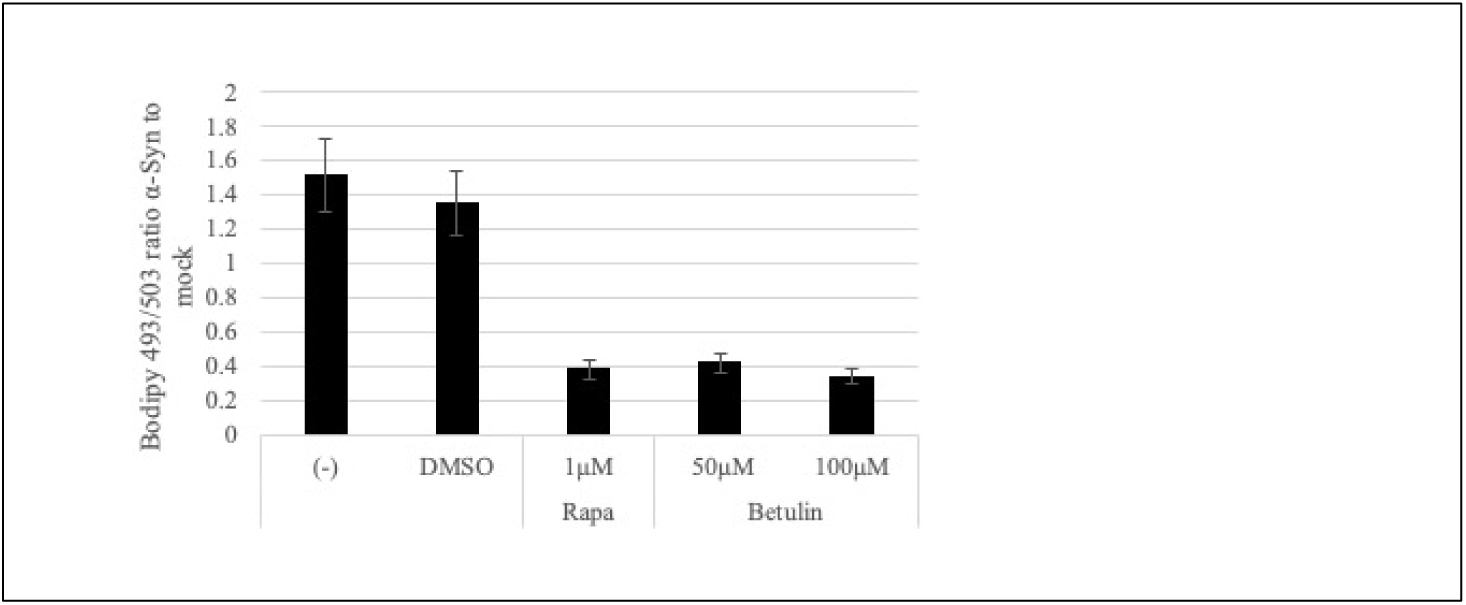
α-Syn -mediated LD- accumulation is sensitive to Betulin, a SREBP inhibitor. HeLa cells expressing α-Syn-mRuby or a mock-mRuby were serum-starved for 16 hours followed by serum-replenishment for 3 hours in the presence of the indicated drug concentrations. Cells were collected and analysed 48 hours from infection by FACS. Mean±SD of n>25000 gated cells.

**S7.**
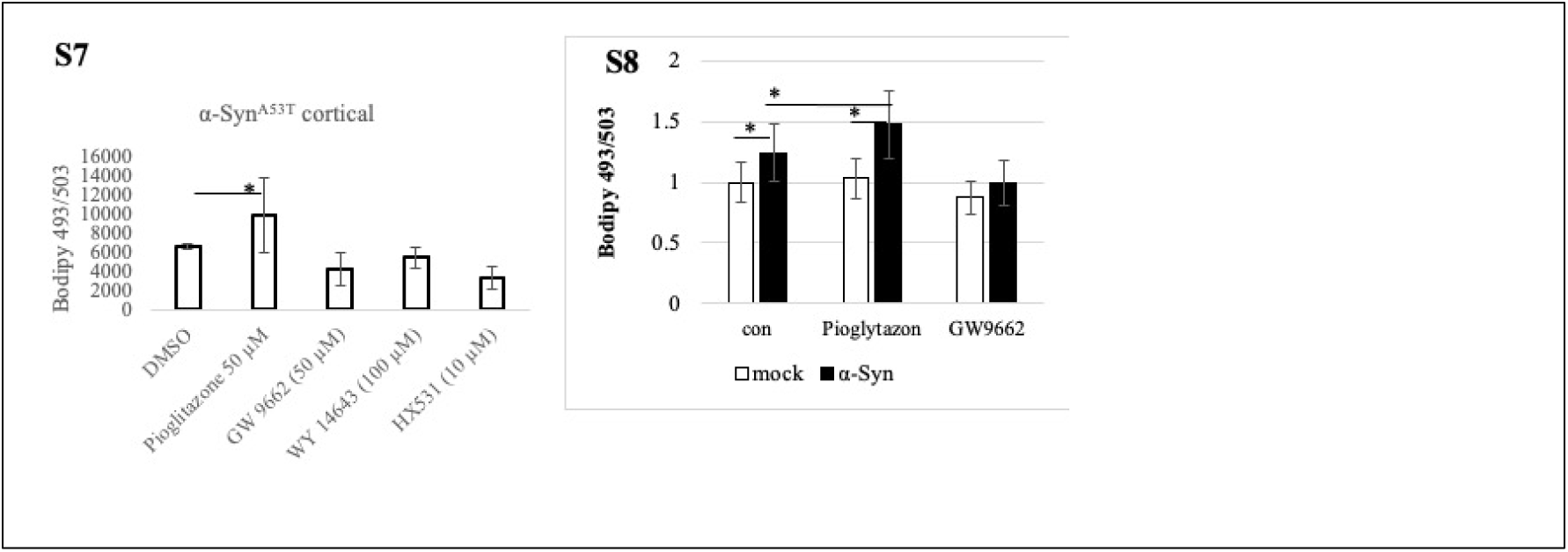
Lipid droplets in primary cortical neurons from α-Syn^A53T^ determined by ICC. Bar graph showing quantitative data of neurons at 10 DIV, treated for 16 hours with GW9662 (50 μM); pioglitazone (50 μM); WY-14643 (100 μM); or HX531 (10 μM). Control cells treated with DMSO. Fixed cells immunoreacted with anti Tuj1 and, and stained with Bodipy 493/503. Images obtained using Nikon spinning disc microscope at 40X magnification. Quantifications using QuPath. Mean±SD of n>40 cells. *, P<0.01 ANOVA.

**S8.** Bodipy 493/503 signal in response to activation of PPARγ detected in HeLa cells by FACS. Hela cells infected to express mock-mRuby or α-Syn -mRuby, treated overnight with GW9662 (50 μM); pioglitazone (50 μM) or vehicle (DMSO) and anlysed for LDs following staining with Bodipy 493/503. Mean±SD on n> 45,000 gated cells, *, P<0.05, one way ANOVA.

**S9.**
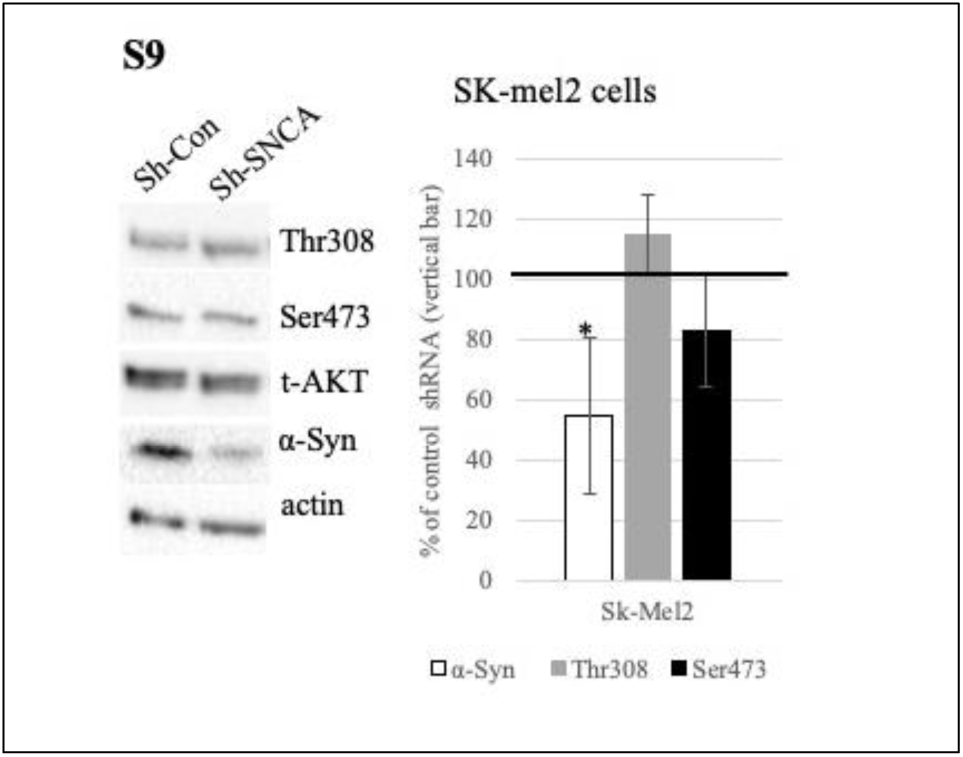
α-Syn expression does not affect AKT activation in SK-Mel2 cells expressing the constitutively active Q61R NRAS mutation. SK-Mel2 cells were infected with either shSNCA or scrambled shRNA control. Seventy-two hours post-infection, AKT activation was assessed. Data are normalized to cells expressing scrambled shRNA (set at 100%, indicated by the vertical line). Bar graph represents the mean ± SD of n = 4 experiments. *, P < 0.01, ttest.

**S10.**
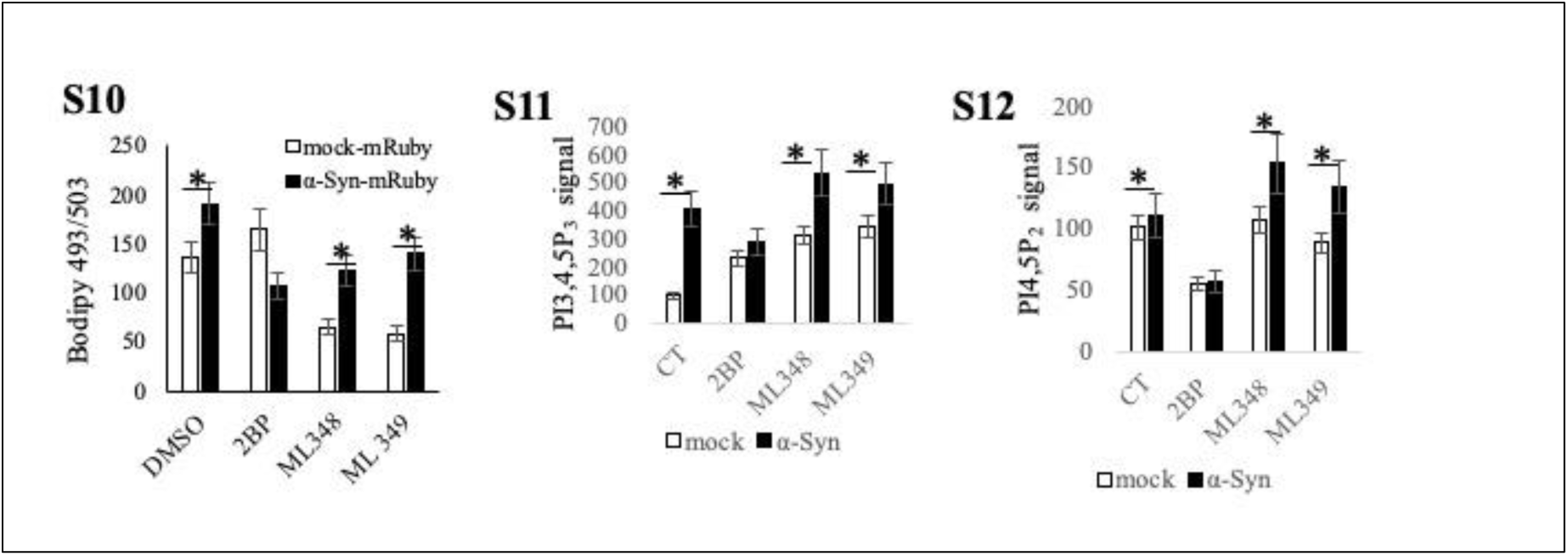
Protein palmitoylation is required for α-Syn to activate PI3K and LDs accumulation. HeLa cells infected with α-Syn-mRuby or mock-mRuby. Sixty hours post infection, the conditioning media was replaced and cells were serum-starved for 14 hours, then, serum (8%) was replenished for 3 additional hours in the presence of the specific inhibitors [ML348 (10 μM), ML349 (10 μM), 2BP (100 nM)]. LDs were determined following staining of live cells with Bodipy 493/503 and analyzed by FACS. Mean±SE of >40,000 gated cells, a representative experiment of 4 repeats * P<0.0001 one way ANOVA. **(S11)** Cells as in (S10) immunoreacted with anti PI3,4,5P_3_ ab (Echelon Biosience). Signal detection by FACS. Mean±SE of n=4 experiments; >40,000 gated cells in each experiment, * P<0.0001. Results presented as % of control cells. **(S12)** Cells as in (S10) immunoreacted with anti PI4,5P_2_ ab. Mean±SE of n=4 experiments; >40,000 gated cells in each experiment, * P<0.0001. Results presented as % of control cells.

